# Homeostatic Plasticity Enables Stable yet Tunable Neuronal Assemblies

**DOI:** 10.64898/2026.05.31.729097

**Authors:** Michelle C Miller, Christoph Miehl, Brent Doiron

**Author notes:** These two authors have contributed equally (Christoph Miehl, Brent Doiron).

## Abstract

Strongly interconnected neuronal populations, called assemblies, dynamically form through synaptic plasticity mechanisms and are thought to be a substrate for memories in the brain. Many models of assembly formation use Hebbian excitatory-to-excitatory plasticity, where coordinated activity strengthens recurrent structure. However, these models typically yield binary assembly outcomes: networks with either weak (no assembly) or maximally strong (assembly) connectivity. We consider networks with a combination of Hebbian excitatory-to-excitatory plasticity and inhibitoryto-excitatory synapses with plasticity that homeostatically stabilizes excitatory neuron firing at a target value. When we set excitatory-to-excitatory plasticity to be homeostatically compliant, in that potentiation and depression are balanced at the homeostatic target firing rate, we find a stable continuum of synaptic strengths, and assembly structure is no longer binary. We use a recurrent network of spiking neuron models and an associated mean-field theory to identify this continuum as a line attractor in synaptic weight space. While along the attractor homeostasis ensures that neuronal firing rates are invariant, the dynamical response properties of the network are quite malleable, with strongly coupled networks having high gain and longer timescale responses. Using our mean-field theory we show how correlated stochastic spiking activity among the excitatory neurons can destroy the line attractor, yet this can be mitigated when correlated inputs are shared across the excitatory and inhibitory neurons. Altogether, we provide a learning framework based on homeostasis, where a tunable and flexible assembly structure is possible.

## I. INTRODUCTION

Groups of neurons with temporally correlated spiking activity are thought to be the basic unit of representations and memory in the brain [1–3]. Evidence suggests that such co-activated groups of cells appear upon stimulus presentation [4, 5], during spontaneous activity [6] and have been linked to animal behavior [7–9]. The physical substrate of this correlated activity is believed to be assembly structure within the network, in which functionally related neurons are connected through strong synaptic connections [3, 10–15]. This long-standing notion goes back to Hebbian theory [16, 17], where long-term synaptic potentiation occurs when pre- and post-synaptic neurons have coincident activation. Indeed, measurements of neural connectivity show that neurons tuned to the same stimuli are both more likely to be connected and share strong connections [18–21].

It is believed that long-term excitatory-to-excitatory (*E*→ *E*) synaptic plasticity underlies assembly formation [3, 22]. A large number of computational studies show how assembly formation can follow from biologically observed plasticity rules [11–13, 23]. However, pure Hebbian *E* →*E* plasticity is generically unstable: recurrent excitation amplifies itself and drives unchecked growth in excitatory neuron firing rates and *E* →*E* synaptic weights [12, 22, 24, 25]. A variety of mechanisms have been used to stabilize assembly formation: weight normalization of the incoming weights into a neuron [24], metaplasticity in which the plasticity rules themselves adapt [26, 27], or upper bounds on synaptic weights [13, 14, 28].

These computational mechanisms can be linked to experimentally described homeostatic and compensatory processes, including synaptic scaling [29, 30] and heterosynaptic plasticity [31, 32]. However, such global homeostatic mechanisms are generally too slow, by themselves, to stabilize fast Hebbian *E* → *E* plasticity in recurrent network models [25, 33, 34]. Experimental and theoretical work point to homeostatic inhibitory to excitatory (*I* → *E*) plasticity as a robust and fast stabilizing mechanism [35–38]. Homeostatic plasticity here refers to modifications of inhibitory synapses that keep each postsynaptic neuron near a *target* firing rate during (and after) learning [25, 34–36, 38]. Such experimentally supported homeostatic *I* → *E* plasticity [39–42] occurs on the timescale of *E* → *E* plasticity [40, 43] and can keep a balance between excitation and inhibition (*EI* balance) [12, 35, 44–46]. In computational models, homeostatic inhibitory plasticity that promotes *E/I* balance in turn stabilizes assembly formation [12, 13, 37].

While homeostatic plasticity can facilitate network firing rate stabilization, when combined in computational models with Hebbian *E* → *E* plasticity rules, there remains a severe limitation in stabilizing the weight dynamics: network models still need a hard [13, 28] or soft [12, 23, 47–49] synaptic weight threshold. This is because after sufficient training, an assembly of strongly coupled neurons continues to potentiate recurrent connections post training through a resurgence of coordinated spiking activity. By contrast, with insufficient training the connections depress and assembly structure dissipates after training [12, 13, 23, 28]. Thus, in the long-time limit post-training dynamics tend toward one of two endpoints: recurrent connections either approach their upper bound, producing a strongly coupled assembly, or weaken until the learned structure dissipates and therefore does not store the training experience [32, 47, 50–52]. This binary nature of assembly structure prevents graded differences in training from being stably retained in the strength of recurrent assembly connectivity and its response properties.

Additionally, assembly learning comprises neuronal allocation and subsequent recurrent synaptic plasticity. Here, we model the latter within a pre-allocated candidate assembly using a recurrent spiking network with homeostatic *I* → *E* synaptic plasticity, and ratedependent Hebbian *E*→ *E* plasticity with one crucial constraint. We introduce the concept of *homeostatically compliant E* → *E* Hebbian plasticity: when a postsynaptic *E* neuron fires at the homeostatic target rate set by the *I* → *E* synaptic plasticity, then the potentiation and depression of the *E* → *E* plasticity are balanced and there is no net weight change (this case was also studied in [37, 53]). We show that this combination of homeostatic *I* → *E* and homeostatically compliant *E* → *E* plasticity can lead to a continuum of stable configurations in synaptic weight space, so that trained assemblies need not be binary in strength (see also [37]). To gain a deeper understanding, we develop a mean-field model of the weight dynamics and describe the continuum as a line attractor in the space of *I* → *E* and *E* → *E* weights. Using our theory we derive the model parameter conditions for the line attractor to exist.

While synaptic line attractors have been described in previous computational work [37, 53], their functional role in assembly learning within recurrent networks remains unstudied. One possibility is that attractor dynamics along a continuum provide a mechanism for assemblies to exhibit graded functional states rather than discrete weak-or-strong connectivity regimes. Neurons have heterogeneous gains, tuning widths and amplitudes that depend on context [54], recent spiking history [55], and behavioral state [56–59]. Since neural assemblies are composed of such heterogeneous neurons, their collective responses may likewise vary in a graded manner, reflecting differences in neuronal participation, activation strength, and response reliability [60]. Consistent with this view, population activity can vary across repeated experiences and across days [61–63], and the strength and timescale of population responses can reflect behavioral variables such as confidence, salience, or memory strength [9, 63]. We use our model and associated mean-field theory to show that for synaptic weights on the line attractor, while the network firing rates are invariant due to homeostasis, the dynamical response properties of neurons in the network can vary substantially.

Finally, the neutral stability inherent along a line attractor often prompts an analysis of how fluctuations affect dynamics [64–67]. Using our linear response theory we show how correlated external fluctuations that cause shared variability across neurons create a positive deterministic drift in synaptic weight space, effectively destroying the line attractor. Fortunately, such drift can be mitigated by coordinating external inputs to excitatory and inhibitory neurons, to allow recurrent network dynamics to cancel shared variability [68–70].

Our findings broaden the functional role of homeostatic plasticity in recurrent networks. While, canonically, the literature has discussed how homeostatic plasticity stabilizes networks through preventing runaway dynamics, we propose it serves an accompanying function of encoding variable and tunable response properties. That is, with homeostasis, when a network learns, it learns a particular response, not just to respond.

## II. RESULTS

### Recurrent excitatory-inhibitory network model with synaptic plasticity

To explore how excitatory and inhibitory learning rules shape emergent synaptic weight dynamics, we construct a recurrent network model with *N*_*E*_ excitatory (*E*) and *N*_*I*_ inhibitory (*I*) spiking neuron models that are each driven by *N*_*X*_ external (*X*) neurons (two external populations) (**Fig. 1a**; see **Methods**). To simplify exposition we take *N*_*E*_ = *N*_*I*_ = *N*_*X*_ ≡ *N*. Throughout, we interpret the modeled E population as a candidate excitatory assembly whose membership has already been allocated. The remainder of the recurrent and feedforward circuits are not modeled explicitly. Their background influence is coarse-grained as statistically stationary external drive common to all neurons. The spiking activity of each *E* and *I* neuron is stochastically sampled according to a Hawkes process [71, 72]. Briefly, the probability of neuron *i* in population *α* ∈ {*E, I*} emitting a spike over a short time interval is proportional to a dynamic intensity function (firing rate) 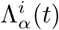, which obeys:

**FIG. 1.**
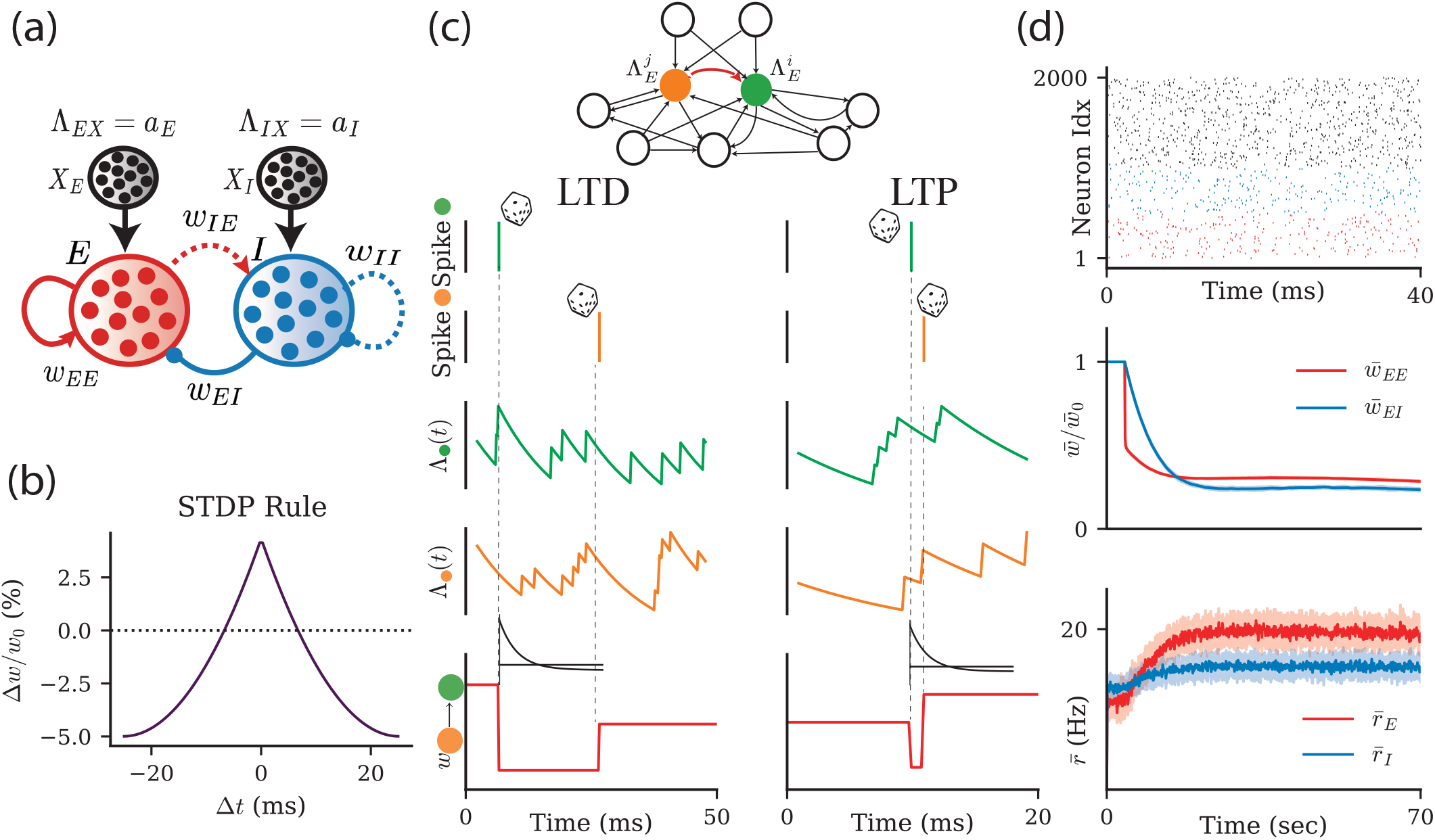
Spiking and STDP framework for recurrent Hawkes process network. **a**. Schematic of the recurrent excitatory-inhibitory (*EI*) network. Neurons in external populations Λ_*αX*_ (*α* ∈{*E, I*}) fire as homogeneous Poisson processes with rates *a*_*E*_ and *a*_*I*_, respectively. In the *EI* network plastic synaptic connections are solid red (*w*_*EE*_) and blue (*w*_*EI*_), while non-plastic connections are dashed (*w*_*IE*_ and *w*_*II*_). **b**. Schematic of the spike timing-dependent plasticity (STDP) as a function of the delay between pre- and postsynaptic spikes Δ*t*. Weights potentiate if pre- and postsynaptic neurons spike with a small delay |Δ*t*|, and depress otherwise. **c**. Top: Schematic of the inputs to *E* neurons *i* and *j* with intensity functions 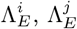 (for *i, j* ∈ {1, …, *N*_*E*}_). We show projecting external input neurons. Red connection indicates the *j → i* weight in the *E* population undergoing plasticity. Bottom: Spike sampling depends probabilistically on the intensity function Λ(*t*). Left: A large time delay Δ*t* between pre- and postsynaptic spikes leads to long-term depression (LTD). Right: A small time delay Δ*t* causes long-term potentiation (LTP). **d**. Top: Spike train raster of excitatory (red), inhibitory (blue), and external (black) neurons. Middle: Mean weight dynamics normalized to initial mean weight strength 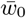 of excitatory 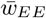 and inhibitory weights 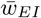. Bottom: Dynamics of mean excitatory 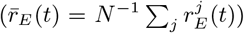 and inhibitory firing rates 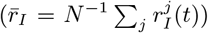. Solid line is an average over 15 realizations of the spike production via 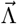 and the shaded area is the standard deviation of the mean.

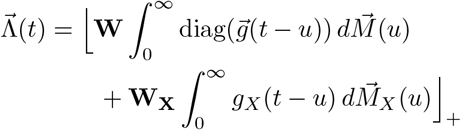

Here 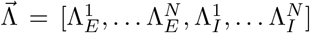 is the vector of intensity functions for the recurrent neurons. We make the assumption that the argument of the rectified linear function is always above threshold. The normalized kernel 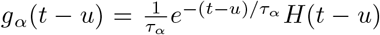 *H*(*t* −*u*) describes the postsynaptic current elicited by a spike from presynaptic population *α* ∈ {*E, I, X*}.*H*(*s*) is a Heaviside function to enforce causality, and 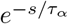 is the postsynaptic ‘current’ for population *α* ∈ {*E, I*} having time constant *τ*_*α*_ (we take *τ*_*E*_ = *τ*_*X*_ and *τ*_*I*_ = 2*τ*_*E*_). Finally, 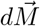 contains the differentials of the *N*_*E*_ + *N*_*I*_ spike count processes, each of which takes the value 1 if a presynaptic neuron spikes over a small interval *du*, and 0 otherwise. Thus, if a neuron spikes at time *u*, the spike will contribute 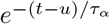 to the intensity functions of other neurons at time *t > u*. A realization of the spike train from neuron *i* in population *α* ∈ {*E, I*} is 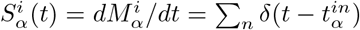, and we define 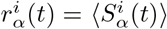, where ⟨*·*⟩ denotes an expectation over realizations of the spike train intensity function 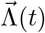.

The matrices **W** and **W**_*X*_ are the 2*N* × 2*N* recurrent and feedforward synaptic couplings, respectively. The recurrent matrix is set as 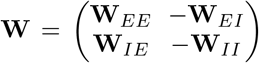, where the individual terms in each block (e.g. **W**_*EE*_) are initialized with entries 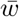 (e.g. 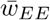). There are two external populations (*X*_*E*_ and *X*_*I*_) which feedforward synapse onto the *E* and *I* populations. We set **W**_*X*_ to have a diagonal block structure (meaning that pairs of *E* neurons receive common inputs, but an *E* and *I* neuron pair does not). The external neurons fire as independent, homogeneous Poisson processes with an intensity function Λ_*αX*_ = *a*_*α*_ ≥ 0 with *α* = {*E, I*}.

In our study, we let the *E*→ *E* and *I* → *E* connections be plastic, while the *E* → *I, I* → *I, X* → *E* and *X* → *I* are non-plastic. We model long-term synaptic plasticity based on a spike-timing-dependent plasticity (STDP) rule, in which long-term potentiation (LTP) or depression (LTD) depends on the relative timing of pre- and postsynaptic spikes. STDP has been described in a variety of studies and has a rich diversity of forms [22, 73, 74]. We follow past work [75–79] and take the synaptic weight 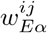 between a presynaptic neuron *j* in population *α* ∈ {*E, I*} and a postsynaptic neuron *i* in the *E* population to evolve according to the STDP rule (see **Methods):**

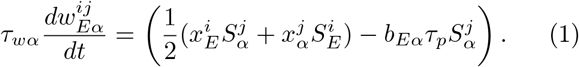

The timescale of weight dynamics is set by the plasticity timescale *τ*_*wα*_. Throughout we take *τ*_*wE*_ = 3*τ*_*wI*_.

LTP recruitment is determined from the temporal overlap between pre- and postsynaptic spiking activity. This overlap depends on the eligibility traces 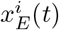 and 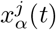 which filter spike trains with an exponential decay (**Fig. 1b**; **Methods**):

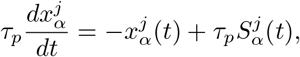

where *τ*_*p*_ is the eligibility timescale. The form of the LTP terms in the STDP dynamics (the *xS* terms in Eq. 1) gives an acausal symmetric Hebbian rule [28, 80, 81]. Specifically, the ordering of pre- and postsynaptic spikes is unimportant, and nearly coincident spiking recruits LTP (**Fig. 1b**). Finally, LTD is recruited by isolated presynaptic activity in population *α* ∈ {*E, I*} and its magnitude is set by the coefficient *b*_*Eα*_.

In total, in our model spikes occur stochastically according to the intensity function 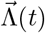, and synaptic weights change based on the relative timing of pre- and postsynaptic spikes (see schematic in **Fig. 1c**). Simulations of our network lead to approximately asynchronous spiking dynamics of recurrent *E* and *I* neurons (**Fig. 1d; top**). If the initial synaptic weights are strong then both the *E* → *E* and the *I* → *E* weights decay to depressed steady state values (**Fig. 1d; middle**). The depression of synaptic weights is mirrored by the *E* and *I* neuron firing rates converging to moderate values (**Fig. 1d; bottom**).

### Homeostatic compliance of excitatory plasticity

We first consider the plasticity of the *I* → *E* connections. The LTD coefficient *b*_*EI*_ sets a desired target *E* population firing rate 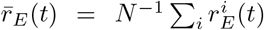. Specifically, the evolution of 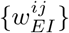 is such that 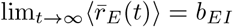 (**Fig. 1d; bottom**; *b*_*EI*_ = 20 Hz). This result is due to the negative feedback loop between the *I* → *E* synaptic weights and the *E* population firing rates. To elaborate, if a postsynaptic neuron *i* in the *E* population has a firing rate 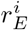 that is above (below) *b*_*EI*_ then Eq. (1) is such that the synaptic weights 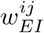 will potentiate (depress) so as to lower (raise) 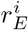. In this way, the *I*→ *E* plastic pathway acts as a homeostatic mechanism for firing rate, as has been extensively studied [12, 13, 23, 35, 37, 82].

While *I* → *E* plasticity is well known to enact homeostasis, the impact that the LTD coefficient *b*_*EE*_ has on the plasticity dynamics of the *E* → *E* pathway is less explored. The *E* → *E* pathway cannot homeostatically regulate network activity, because it supports a positive feedback loop between potentiating synaptic weights 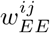 and growing firing rates 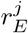 and 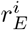. Nevertheless, the form of Eq. (1) for the *E* → *E* pathway mirrors that of the *I* → *E* pathway. The central goal of our study is to explore the plasticity/network activity dynamics for the symmetric STDP case where *b*_*EI*_ = *b*_*EE*_ ≡ *b*. This has been shown in previous work to be necessary for stable learning in feedforward networks [37]. We consider this necessity in recurrent networks.

As a first exploration we test how different initial conditions (*t* = 0) of the mean excitatory weight 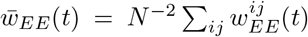 affect the evolution of the network/synaptic dynamics. We find that 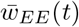 asymptotes to a steady-state value that depends on the initial conditions (**Fig. 2a**). Further, since the *I* → *E* pathway aims to keep the excitatory firing rates at a target value (*b*), a higher initial mean excitatory weight, 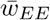, is balanced by a higher mean inhibitory weight, 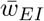 (**Fig. 2b**). In total the network with both *E* → *E* and *I* →*E* plasticity shows a continuum of stable states, often termed a ‘line attractor’, in the 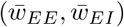 phase space (**Fig. 2c**, dashed line). The homeostatic *I* → *E* plasticity enforces the *E* neuron target firing rate to be 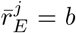 for all synaptic weight initializations, and consequently *I* neurons also converge to a fixed rate along the attractor (**Fig. 2d-e**). In the above example the *E* → *E* pathway satisfies the constraint *b*_*EI*_ = *b*_*EE*_ = *b*, because then 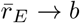 for a range of stable 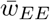 values. Because of this, we say that the excitatory plasticity is *homeostatically compliant* when *b*_*EI*_ = *b*_*EE*_.

**FIG. 2.**
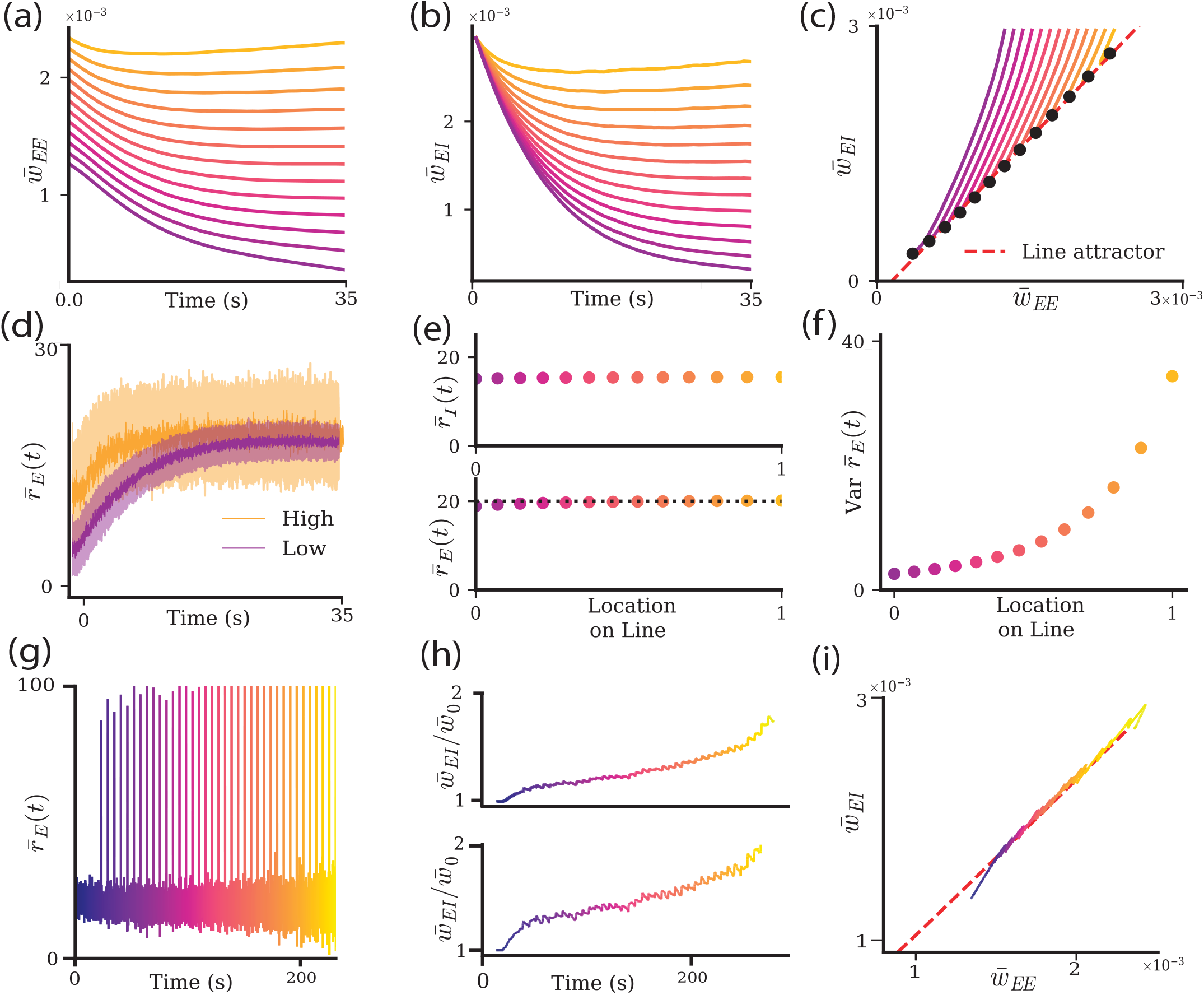
Line attractor in weight space in an *EI* spiking network. **a**. Mean *E → E* weight, 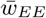, evolution for different initializations (indicated by the line color). **b**. Mean *I → E* weight, 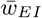, evolution over time for the different 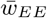 initializations in panel a. **c**. Weight dynamics in the 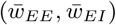 phase space. Colors are as in panel a. Black dots represent the weight after 20 sec of simulation, and the red dashed line (the ‘line attractor’) is a linear regression fit through the black points (*r*^2^ = 0.99). **d**. Mean excitatory firing rate 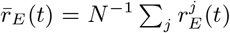 over time, averaged over 5 trials, for two initial conditions from panel a (*b*_*EI*_ = 20Hz). **e**. Mean inhibitory 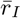 (top) and excitatory rates 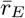 (bottom) averaged over 12-17 sec of simulation time. Location on the line attractor is parametrized from 0 (lowest steady state weights) to 1 (highest steady state weights) in panel c. Dashed line indicates rate set-point *b*_*EI*_ = 20Hz. **f**. The variance of the mean excitatory firing rates, 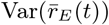 as a function of location on the line attractor (*T* = 5*s* in calculating 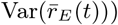. **g**. Mean excitatory rate 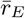 response to the series of pulses over time. **h**. Mean normalized synaptic weights 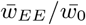 (bottom) and 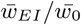 (top) dynamics during the learning paradigm. **i**. 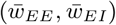 phase space during the learning paradigm.

Do line attractor dynamics for the evolution of synaptic weights require that *E* → *E* plasticity be home-ostatically compliant with the *I* → *E* plasticity? It is possible to create a stable continuum of 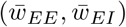 when the *E* → *E* plasticity is a pairwise anti-symmetric rule (causal) such that LTP and LTD are exactly balanced for all (asynchronous) pre- and postsynaptic activity (**Supplement Fig. 1a**). While *b*_*EE*_ = 0≠ *b*_*EI*_, in this case 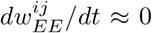 and there is no net evolution of 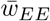 regardless of the *E* population firing rate. While this forms a line attractor in the joint 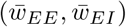 space, it does so in a trivial sense: 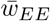 remains fixed at its initial value, and only 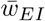 evolves toward the continuum. Conversely, if *E* → *E* LTP and LTD are imbalanced for all (asynchronous) activity in the asymmetric window, then 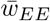 drifts deterministically, and recurrent excitation cannot settle even if inhibition 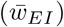 has begun approaching a steady state (**Supplement Fig. 1b**).

Networks employing a more realistic triplet STDP plasticity rule [83, 84] also exhibit *E* firing rate homeostasis with line attractor dynamics (**Supplement Fig. 1c**). Although the triplet rule removes the need to fine-tune the balance between LTP and LTD, it still requires the excitatory and inhibitory plasticity thresholds to agree to support stable sustained dynamics; mismatched thresholds do not (e.g. *b*_*EE*_ = 30 and *b*_*EI*_ = 20; **Supplement Fig. 1d**). We limit our excitatory plasticity rule to a symmetric pairwise rule, particularly for mathematical simplicity, but many of our mean-field stability results below extend to the asymmetric triplet rule.

The *b*_*EE*_ = *b*_*EI*_ constraint can be loosened if the *b*_*Eα*_ terms are themselves allowed to be dynamic [37] (**Supplement Fig. 1e**). In that case, each *b*_*Eα*_ has a different initial value, but each acts as a low-pass filter of the postsynaptic firing rate 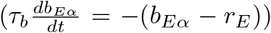. The limiting factor for stability becomes the timescale on which the plasticity thresholds change, *τ*_*b*_. In sum, when *E* → *E* plasticity is homeostatically compliant with the *I* → *E* plasticity, then 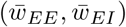 exhibits line-attractor dynamics.

A key observation of our study is that while firing rates are invariant along the line attractor, the dynamic properties of population activity are not. To illustrate, we first consider the stationary variance of the *E* population activity, 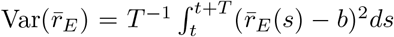, where the domain of integration (*t, t* + *T*) is when the synaptic weights have converged onto the line attractor (so that 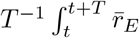 (*s*)*ds* ≈ *b*) where *T* = 5sec. Networks with large synaptic weights 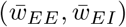 have much larger 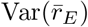 than those with small weights (**Fig. 2f**). This suggests that a network’s intrinsic variability or variability in response to input varies along the synaptic weight line attractor, even while the firing rates 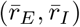 remain invariant.

Having established a continuum of stable synaptic configurations, we next ask how coordinated stimulation of the population impacts the network on the line. We use a stimulation protocol in which each neuron in the *E* population is stimulated with brief (30 ms) input arriving periodically at a rate of 0.2Hz (**Fig. 2g**). We find that this repeated stimulation can induce assembly learning (growth of 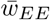) in the network. The mean *E* neuron firing rate responds to each pulse (**Fig. 2g**), and over repeated pulses, 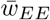 and 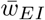 increase in strength (**Fig. 2h**). The synaptic weight increases closely follow the line attractor (**Fig. 2i**) so that the weight changes act like a ratchet, and over the protocol, weight increases build upon past increases. Repeated stimulation of a population can potentiate 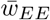 and give rise to a strong, stable assembly.

Taken together, these results show that *E* → *E* and *I* → *E* plasticity does more than prevent runaway excitation through homeostasis. When *E* → *E* plasticity is homeostatically compliant, then a line attractor in 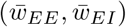 space is produced. While by design the mean firing rates are invariant along this line attractor, the variability grows. Thus, the synaptic line attractor provides a stable yet tunable substrate on which experience can cumulatively strengthen an assembly while preserving its homeostatically regulated mean firing rate. However, this raises the question of whether movement along the synaptic line attractor alters the dynamical stability and functional response properties of the assembly.

### Mean-field model of the synaptic weight line attractor

To gain a more rigorous description of the line attractor in synaptic weight space and understand how it shapes population dynamics, we derive and analyze a mean-field model of the spiking/synaptic network activity. We first assume that the network is in an asynchronous regime, and that the dynamics of 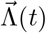 are much faster than the evolution of the synaptic weights **W**(*t*) (i.e *τ*_*r*_ ≪ *τ*_*wα*_).

On the fast timescale of network activity we derive mean-field equations for the population and trialaveraged activity (**Fig. 3a**):

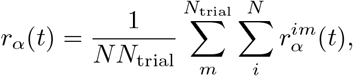

for large *N* and *N*_trial_. For the Hawkes process network model, the mean firing rates of the excitatory and inhibitory populations, 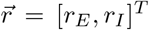, obey the following linear system (**Appendix A**):

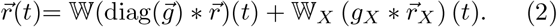

**FIG. 3.**
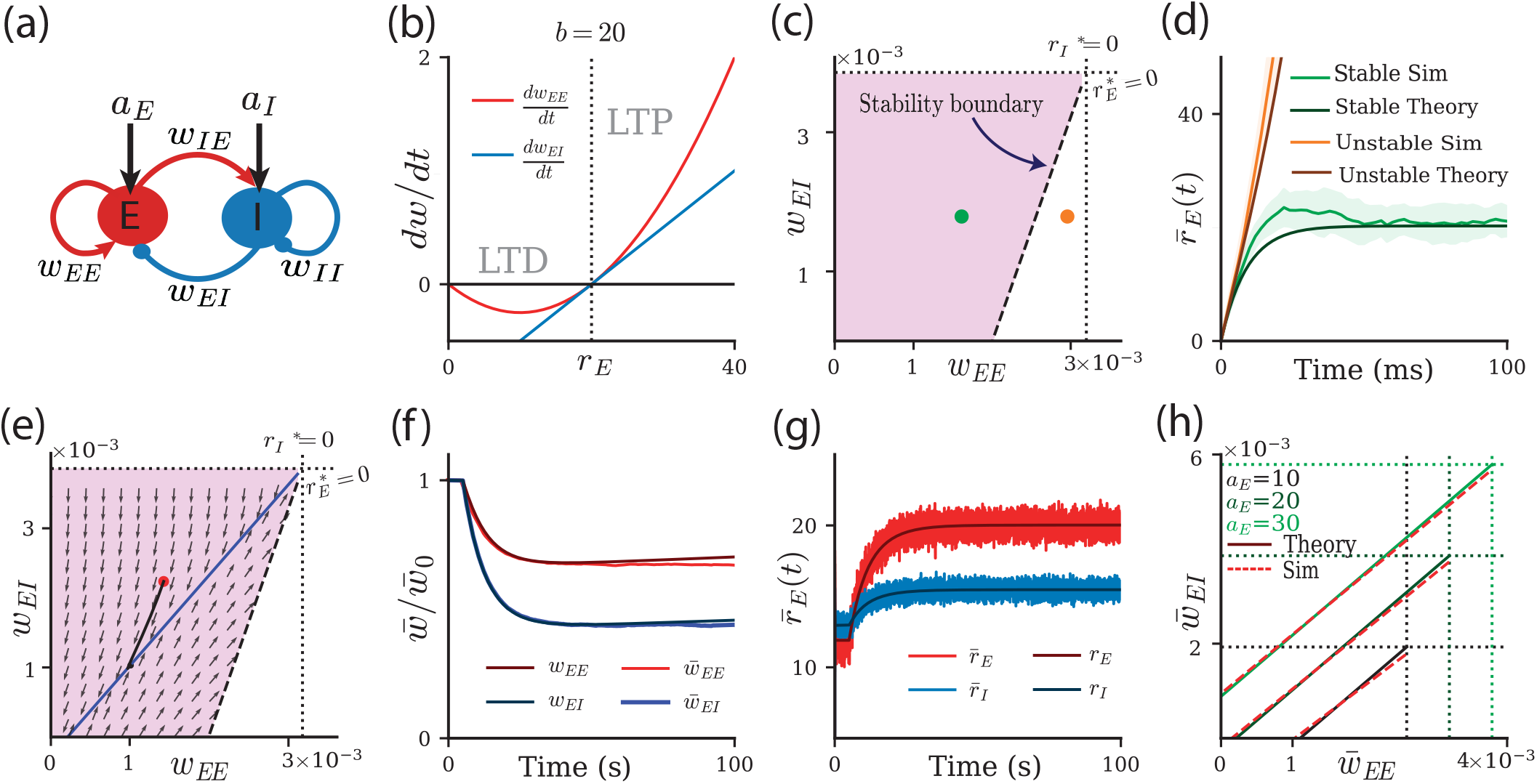
Synaptic weight line attractor in a mean-field model. **a**. Schematic of the mean-field model. An excitatory (*E*) and inhibitory (*I*) population are coupled via synaptic weights *w*_*αβ*_ and receive external input via *a*_*E*_ and *a*_*I*_. **b**. Schematic for the *E →E* (red) and *I →E* (blue) rate-dependent plasticity rule. Weights potentiate for excitatory firing rates *r*_*E*_ above *b* = 20*Hz* (LTP) while depressing otherwise (LTD). **c**. The region of stable rate dynamics is framed from below by 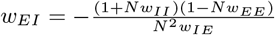 (black dashed line), and from the right by 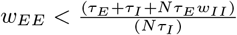 (not shown). From the top and the right, the space is constrained by boundaries for which firing rates would become negative (dotted black lines). For 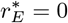, we have the boundary 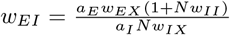 and for *r*_*I*_ * = 0, the boundary is 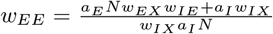. The collection of these boundaries defines a region of stable firing rates, which is indicated by the pink trapezoid in (*w*_*EE*_, *w*_*EI*_) space. The green and orange dots are networks that exhibit stable and unstable dynamics, respectively. **d**. *E* firing rates of the mean-field model (dark colors) and the Hawkes process network model (light colors) for the two weight pairings indicated in panel c (stable is green; unstable is orange). Firing rates from the Hawkes process network model are averaged across the population over 20 trials. **e**. Same as panel c, overlaid with the flow field of *w*_*EI*_ and *w*_*EE*_ weight dynamics. Blue line indicates the line attractor, black solid line is an example trajectory which started from the red dot. **f**. *E → E* 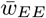 (light red) and *I → E* 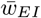 (light blue) weight dynamics over time for the simulation. Dark red and blue are for the mean-field synaptic weights. **g**. Excitatory 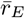 (light red) and inhibitory 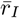 (light blue) firing rate dynamics over time for the simulation. Dark red and blue are for the mean-field firing rates. **h**. Line attractor for different external drives onto the *E* population (*a*_*E*_ = 10, 20, 30Hz). Green lines are from the mean-field model, and red dashed lines from the Hawkes process network model (linear fits, *r*^2^ values are .995, .9998 and .9998). Dotted green lines indicate bounds of zero firing rates.

The normalized causal filters *g*_*E*_, *g*_*I*_, *g*_*X*_ describe synaptic input from the corresponding presynaptic populations, with time constants *τ*_*E*_, *τ*_*I*_, *τ*_*X*_, respectively. See **Appendix A** for the corresponding mean-field ODE. Also,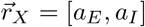 are the external drives onto each population. To distinguish between numerical and meanfield estimates of population activity, we will continue to use the ‘bar’ notation, 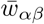 and 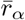, to indicate connectivity and activity calculated from numerical simulations of the Hawkes process model, while *w*_*αβ*_ and *r*_*α*_ will denote solutions from our mean-field theory. Further, we use the notation that bolded matrices (e.g. **W**) correspond to the full-dimensional network, whereas script matrices (e.g. W) denote the reduced mean-field network.

On the slower learning timescale we follow previous studies [76, 79] and derive evolution equations for the mean synaptic plasticity dynamics (**Appendix B**) as:

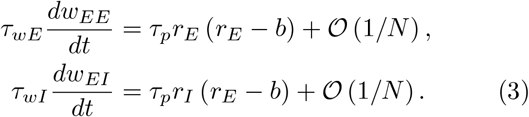

Here the O(1*/N*) term corresponds to the role pairwise spike time correlations would play on the plasticity rules, which for large *N* we can consider negligible in the current network setup. If the firing rate of the excitatory population *r*_*E*_ is above the threshold *b*, the mean *E* → *E* weight *w*_*EE*_ and the mean *I* → *E* weight *w*_*EI*_ will potentiate (LTP), while the weights depress (LTD) otherwise (**Fig. 3b**). Since for each plasticity rule the weight change is multiplied by the firing rate of the respective population, the *I* → *E* plasticity rule has a linear dependency on *r*_*E*_, while the *E*→ *E* rule has a quadratic dependency. Similar dependencies in which the sign and magnitude of *E* → *E* plasticity depend on excitatory firing rates have been found in experimental studies [85–87].

We start by investigating the dynamics of the fast system. The separation of timescales allows us to assume that synaptic weights are effectively fixed as firing rates reach their steady states (**Appendix C 1**). Solving for *r*_*E*_ and *r*_*I*_ at equilibrium from Eq. 2 yields:

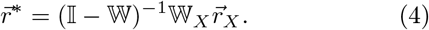

Eq. (4) is only a valid solution for matrices W that permit a stable steady state firing rate 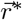. We identify the region in weight space (*w*_*EE*_, *w*_*EI*_) where the firing rate dynamics are stable **(Fig. 3c; pink region)**. We use the stable/unstable characterization to validate that our mean-field model is an accurate quantitative description of the spiking dynamics (**Fig. 3d**; compare dark orange/green to light orange/green traces).

To study the slow synaptic weight dynamics, we consider equilibrium solutions for *w*_*Eα*_ (i.e *dw*_*Eα*_*/dt* = 0 in Eq. 3), and we assume that the fast system (firing rates) is in the stable regime and has reached steady state. In this case the *I* → *E* pathway enacts homeostatic control such that 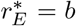. We find that stable weight configurations are described in the large *N* limit by (**Appendix C 2**):

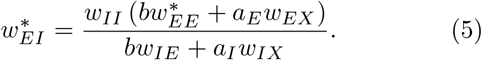

Since 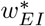 depends linearly on 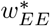, this corresponds to a line attractor in (*w*_*EE*_, *w*_*EI*_) weight space, similar to that observed in the Hawkes process spiking model (**Fig. 2c**). Our mean-field approach reveals how the slope and intercept of the line attractor depend on the (non-plastic) *I* → *I* and *E*→ *I* weight strengths (*w*_*II*_ and *w*_*IE*_), *E* and *I* input parameters (*a*_*E*_*w*_*EX*_ and *a*_*I*_*w*_*IX*_) and the plasticity threshold *b*. The system in Eq. (3) defines a flow field (**Fig. 3e**) and, similar to the Hawkes process model, the mean synaptic weights evolve to a steady state (**Fig. 3f**) which sits along the line attractor. Further, while on the line attractor, the *E* population has a mean firing rate of *b* = 20 Hz (**Fig. 3g**). We confirm that the mean-field estimate of the line attractor matches the line attractor found in the Hawkes process model by changing one example parameter (the external drive onto excitatory neurons, *a*_*E*_). Increases in *a*_*E*_ result in a larger regime of stable firing rates, resulting in an upward shift of the line attractor, like in the Hawkes process model (**Fig. 3h**).

### Conditions for line attractor dynamics

The full system of dynamic firing rates and synaptic weights can co-evolve so that *r*_*E*_ → *b* and (*w*_*EE*_, *w*_*EI*_) converge onto the line attractor; however, this is not guaranteed for all network and plasticity parameters. We require two conditions to be met. First, the line of fixed points must be stable (i.e., a line attractor instead of a line repeller). Second, the initial conditions of (*w*_*EE*_, *w*_*EI*_) must be in the basin of attraction of the line attractor.

#### Stability

Using linear stability analysis, we identify two regimes in which the line of fixed points is an attractor (**Appendix D**, see also [53]). First, stability is guaranteed if we satisfy (in the large *N* limit) the condition:

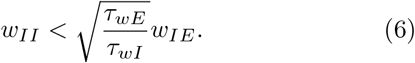

It should be noted here that choosing faster inhibitory than excitatory plasticity dynamics, as is often assumed in previous studies (e.g. [37, 88]), as well as in our case where *τ*_*wE*_*/τ*_*wI*_ = 3, can ensure stable dynamics, though it is not a necessary condition. Here, we see this condition constrains two properties of the network: 1) timescales of the plasticity rules, and 2) circuit properties through weak or strong inhibitory feedback. For example, if inhibitory plasticity were slow, the circuit would need to maintain strong *E* → *I* feedback or weak *I*→ *I* inhibition to remain stable. If Condition (6) is not met, line attractor dynamics can still occur if the plasticity threshold *b* fulfills a second inequality (in the large *N* limit):

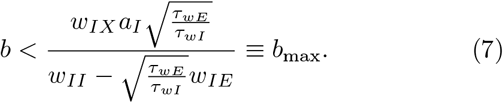

Condition (7) defines the maximum threshold *b*_max_ that supports line attractor dynamics, which was unbounded when Condition (6) was met.

We can visualize these two conditions in (*w*_*IE*_, *w*_*II*_) parameter space (**Fig. 4a**). The grey region represents when the line of fixed points is always an attractor for any *b* (regime 1), while the colored region admits line attractor dynamics for *b < b*_max_ (regime 2). The red, pink, and purple dots represent three different choices of *w*_*IE*_ used in subsequent panels. In regime 1, Condition (6) is satisfied, so *b*_max_ is unbounded, denoted by the blue shaded region (**Fig. 4b**). In regime 2, Condition (7) must be met, and *b*_max_ rapidly decreases as *w*_*IE*_ decreases (orange shaded region) (**Fig. 4b**). Recall that the plasticity threshold *b* is the homeostatic setpoint of the *E* firing rate, hence it defines the firing rate of the *E* population on the line attractor 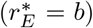. Thus, while in regime 2, even if dynamics are stable, steady state firing rates will be constrained. In total, the non-plastic weights *w*_*IE*_ and *w*_*II*_ together with the plasticity timescales define whether the line of fixed points is an attractor (and not a repeller).

**FIG. 4.**
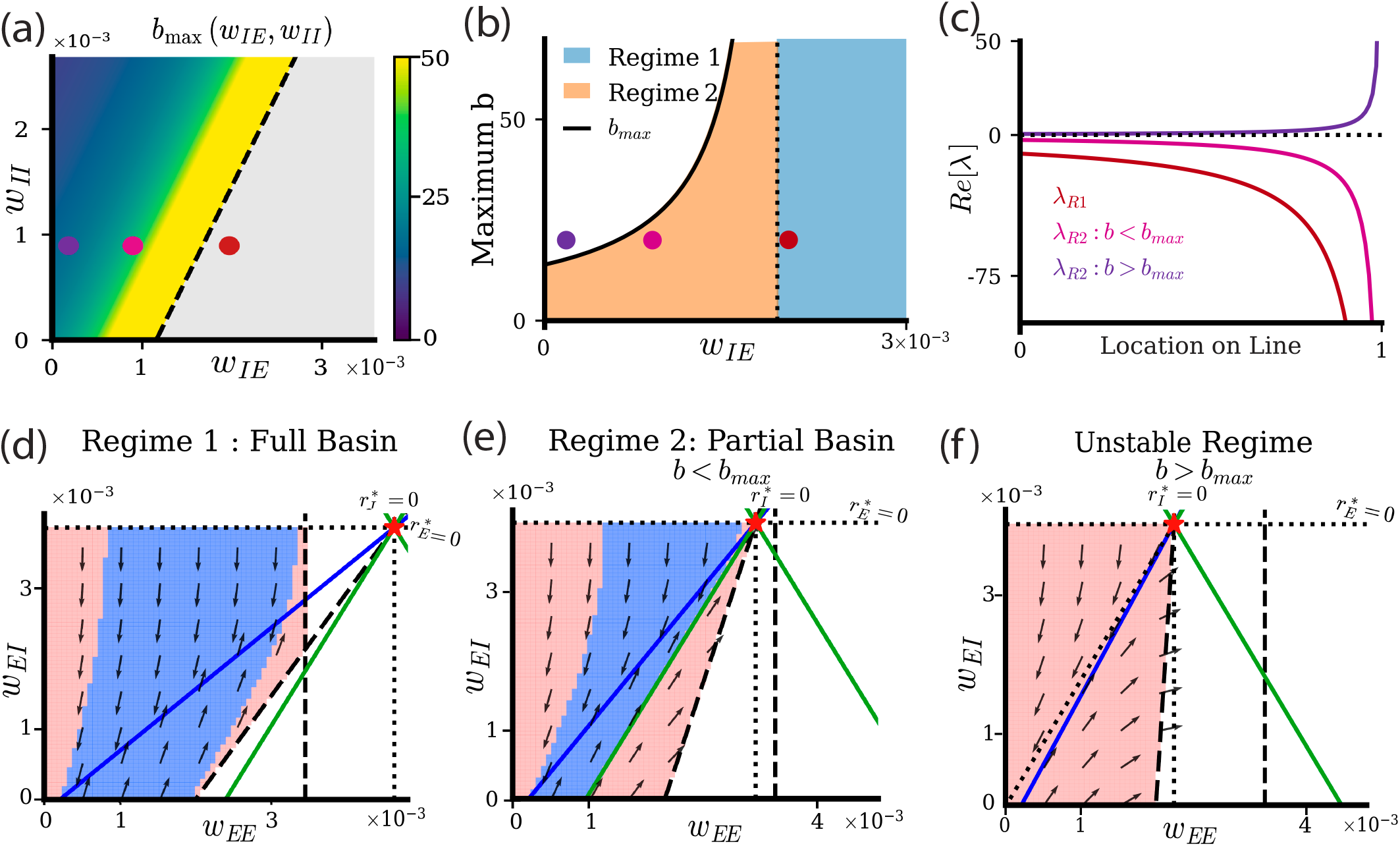
Basin of attraction of the line attractor in the mean-field model. **a**. The upper bound on the plasticity threshold, *b*_max_ for the line of fixed points to be an attractor. The grey region denotes there is no upper bound (regime 1). Yellow to blue color indicates the finite and decreasing *b*_max_ (regime 2). Red, pink, and purple dots indicate parameter choices in subsequent panels. **b**. *b*_max_ as a function of the non-plastic *E → I* weight (*w*_*IE*_) at fixed *w*_*II*_. **c**. The real part of the nonzero eigenvalue of the weight dynamics Re[*λ*] as a function of the location along the line of fixed points. Colors indicate regimes from panel a. **d**. Flow field of *w*_*EE*_ and *w*_*EI*_ weight dynamics in regime 1. Blue line indicates the line attractor, black dotted lines indicate 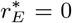 and 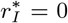, black dashed line separates stable from unstable firing rate dynamics, and the green line is the stable manifold of the non-hyperbolic fixed point (red star at 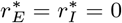). Blue shaded areas are basins of attraction for the line attractor, and red shaded areas are regions where weights do not converge to the line attractor. **e**. Same as d for lower value of *w*_*IE*_ in regime 2, and *b < b*_*max*_. **f**. Same as d and e for a lower value of *w*_*IE*_ in regime 2, and *b > b*_max_.

The exact location on the line attractor, (*w*_*EE*_, *w*_*EI*_) determines the degree of linear stability. We quantify this by calculating the eigenvalues of the slow system (**Appendix D**). For a line attractor, one eigenvalue is always zero and the other non-zero. If the real part of the non-zero eigenvalue (*λ*_nz_) is negative, the dynamics are stable; and unstable otherwise (**Fig. 4c**). The red, pink, and purple curves represent the red dot (stable in regime 1), magenta dot (stable in regime 2) and purple dot (unstable in regime 2) (**Fig. 4a,b**). Specifically, if the system is in one of the stable regimes that fulfill either Condition 6 (red line) or Condition 7 (pink line), the real part of *λ*_nz_ is negative and decreases as a function of location on the line attractor. When choosing *b > b*_max_ (purple line), the attractor becomes a repeller, and the slow system is unstable. It’s notable that while it appears the slow system becomes more stable high up on the line (e.g. the eigenvalues become more negative), suggesting more stable dynamics, the fast system is becoming increasingly unstable. This drives the eventual unstable dynamics at the rate stability boundaries (as well as a breakdown in fast-slow decomposition).

#### Basin of attraction

To gain a deeper understanding of how the line attractor changes its properties, we investigate the dependence of the line attractor and its flow fields for decreasing values of *w*_*IE*_, effectively moving across the different stability conditions (**Fig. 4a**; red, pink and purple dots). Decreasing *w*_*IE*_ increases the slope of the line attractor following Eq. (5), consequently reducing the area of attractor dynamics in (*w*_*EE*_, *w*_*EI*_) space (**Fig. 4d-f**). The area of stable rate dynamics can be further separated into two regions: a basin of attraction where weight dynamics will converge to the line of fixed points (**Fig. 4d**; blue area) and a region where weight dynamics do not converge to the line attractor (**Fig. 4d**; red area).

In regime 1 (**Fig. 4a**; red dot), the basin of attraction is bounded to the right by the firing rate stability boundary (black dashed line) and from above by the 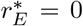 line (black dotted line). To the left, it is bounded by trajectories that ultimately converge to *w*_*EE*_ = 0. Some trajectories within the trapezoidal region to the right of the line attractor do not necessarily converge on the line. This is likely due to a breakdown of timescale separation as the blue diagonal and black vertical dashed lines are firing rate stability boundaries. Decreasing *w*_*IE*_ into regime 2 shrinks the basin of attraction. It is now additionally bounded to the right by a separatrix (**Fig. 4e**; green line), where trajectories to the left of the separatrix converge to the line attractor, while trajectories to the right diverge away from the attractor. Decreasing *w*_*IE*_ further into the unstable regime (**Fig. 4a**; purple dot) removes the basin of attraction entirely because the separatrix moves past the line attractor, leading to a fully unstable line of fixed points, a line ‘repeller’ (**Fig. 4f**; see **Appendix E** for details).

The above conditions in equations 6 and 7 on the plasticity threshold also hold if we use *I* → *E* and *E* → *E* plasticity rules that are a cubic (as opposed to the current quadratic form) function of the postsynaptic firing rates. This has been studied in the context of mean-field dynamics for the triplet rule [37, 89], in which triplets of spikes determine the evolution of synaptic potentiation. The constraints on the plasticity threshold hold given these rules, suggesting generalizability of the linear attractor stability for homeostatic rules (**Appendix F** for details).

### Calculating the network response properties in the mean-field model

Having established when homeostatically compliant plasticity produces a stable line attractor, we next ask how synaptic configurations with identical steady-state firing rates functionally differ. Although positions along the attractor are indistinguishable in their baseline population firing rates, they contain different recurrent excitatory and inhibitory coupling strengths and may therefore respond to transient inputs differently.

To begin, we perform perturbations of the steady state *E* population for networks with synaptic weights along the line attractor (**Fig. 5a**). For (*w*_*EE*_, *w*_*EI*_) pairings that are low on the attractor we observe small and shortlived responses (**Fig. 5b**), while for (*w*_*EE*_, *w*_*EI*_) pairings that are high on the attractor there are large and longlived responses (**Fig. 5c**). Our mean-field model well describes the transient population rate dynamics from the Hawkes process model (**Fig. 5b**,**c**; compare dark colored lines and light colored lines). We aim to use our mean-field theory to give mechanistic insight into these results.

**FIG. 5.**
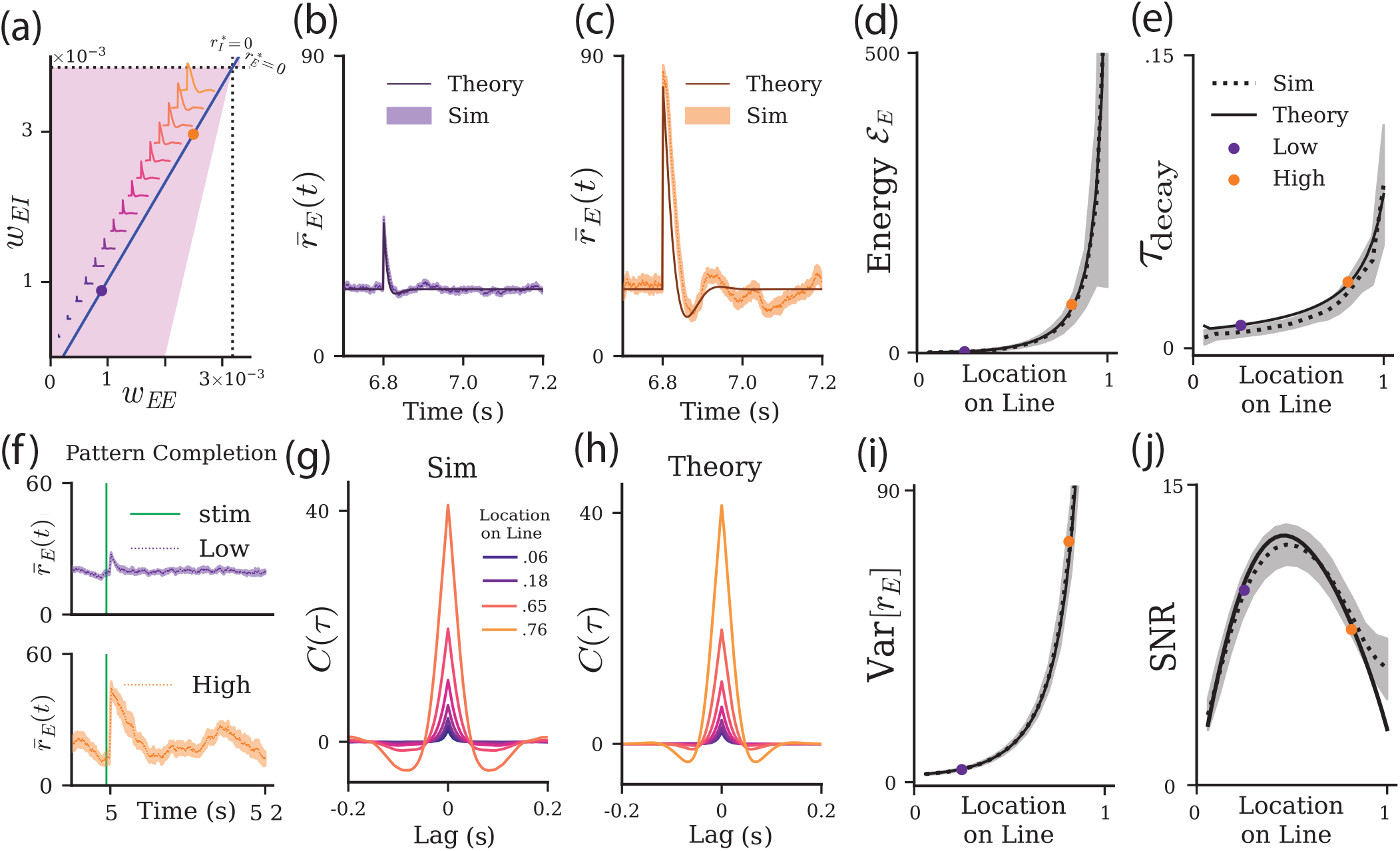
Network response properties vary along the synaptic weight line attractor in the mean-field model. **a**. Two example networks with different initial conditions along the line attractor, one low (purple dot) and one high (orange). Traces of *E* population response to a perturbation are plotted along the line for corresponding coordinates on the line attractor. **b**. Upon receiving a perturbation, the network with weaker coupling has a short, rapid response. Dark purple is theory, light purple is simulations of the Hawkes process network. **c**. The network with stronger recurrent coupling has a stronger rebound in response. Dark orange is theory. Light orange is simulation. **d**. We parametrize location on the line attractor varying from 0 to 1 starting with where the line attractor crosses *w*_*EE*_ = 0 to the coordinate 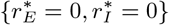. The energy, ε _*E*_, grows along the line attractor for theory (black solid) and simulations (black dotted). **e**. The relaxation time to a perturbation, *τ*_decay_, grows along the line attractor for theory (black solid) and simulations (black dotted). **f**. Mean excitatory rate 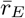 for network low (top, purple) and high (bottom, orange) on the line when only 20% of the network is stimulated. **g**. Autocorrelation function of the mean activity of the E population in simulation for different positions on the line. **h**. Same as g but for theory. **i**. The trial-to-trial variability, Var[*r*_*E*_], grows nonlinearly along the line attractor for theory (black solid) and simulations (black dotted). **j**. The signal-to-noise ratio varies non-monotonically on the line attractor.

Consider the implicit solution for our mean-field network activity:

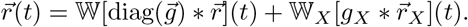

Moving to Fourier space and solving for 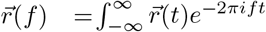 *dt* yields:

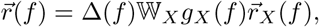

where we have introduced the frequency-dependent propagator:

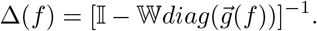

The propagator defines how activity reverberates through the network after being filtered by synaptic dynamics 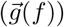. It is the central object governing both response dynamics and variability [90–94]. Once Δ(*f*) is known, (i) network stability, (ii) perturbation response properties, (iii) dynamical and trial-to-trial variability of population responses, and (iv) signal-to-noise ratio can all be expressed in terms of Δ(*f*) or its eigenvalues. Historically, linear response theory has used the propagator to understand these properties in networks of fixed connectivity [90, 91, 93]. We will focus our study on how different synaptic weight pairings (*w*_*EE*_, *w*_*EI*_) along the line attractor determine the propagator Δ(*f*), and hence affect the dynamical response properties of the *EI* network (see **Methods**).

#### Network equilibrium and stability

We first remark that the zero frequency (*f* = 0) mode gives us the equilibrium firing rates in terms of the propagator:

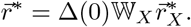

For (*w*_*EE*_, *w*_*EI*_) pairings that are on the weight line attractor, the firing rates 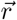 are invariant (**Fig. 2**), making 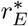 invariant (and equal to *b* when on the synaptic weight line attractor).

The stability of 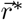 can also be understood through the propagator Δ(*f*). As *w*_*EE*_ and *w*_*EI*_ grow so does Δ(*f*), meaning the reverberations throughout the network become stronger, and compromise stability of 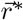. Indeed, as Δ(*f*) grows, an eigenvalue of 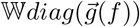 approaches 1, after which the rate dynamics become unstable since 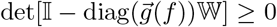. We can obtain the same stability conditions for the fast system with the propagator as those found from traditional linear stability analysis (**Appendix G 1**).

#### Dynamical response

We next consider how the circuit responds to a perturbation of the E population activity. We compare and contrast a network with synaptic weights 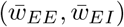 at the bottom of the line attractor to a network with synaptic weights at the top of the line attractor (purple and orange dots in **Fig. 5a**). Once synaptic weight and rate dynamics reach steady state values, we perturb the network by making each *E* neuron spike, causing every *E* neuron synapse to be activated (at time *t*_*p*_). Even though the steady state mean *E* neuron firing rates are identical for the two networks (at the homeostatic target rate *b*), we find they nevertheless show distinct dynamical response properties. The network with weights lower on the line attractor responds to the perturbation with a weak increase in activity, which rapidly dampens (**Fig. 5b**; purple trace). By contrast, the network with synaptic weights high on the line attractor shows an amplified and prolonged dynamical response (**Fig. 5c; orange trace**).

We reproduce these perturbations in the mean-field model (**Fig. 5b/c; dark purple/orange traces**) and likewise use the theory to quantify the differences in the network response. Perturbing around the equilibrium solution gives a closed-form expression for the response to an injected pulse 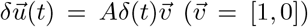 in which *A* is the amplitude of the perturbation. The resulting rate perturbation is 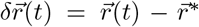. The time evolution of a perturbation depends on the propagator 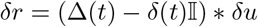 (**Appendix G 2**).

We can use these perturbation evolution equations to calculate the time-integrated squared response (or the ‘energy’) for different points along the line attractor: 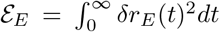 (see **Appendix G 3**). This measures the size of the perturbation over time as it quantifies the sensitivity of the network’s dynamic response.

We find that 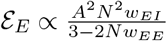, and when written in terms of the time domain propagator we have:

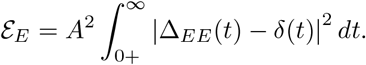

The energy *E*_*E*_ grows as (*w*_*EE*_, *w*_*EI*_) moves upward along the line attractor (**Fig. 5d)**. By integrating over the propagator, *E*_*E*_ quantifies the total reverberatory strength of the perturbation in the excitatory population. We are able to reproduce this relationship by measuring the energy from the simulation traces: 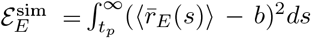. This reinforces the notion that synaptic pairings 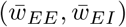 high on the line attractor produce networks with dynamics that are more sensitive to input perturbations than networks with pairings 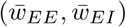 low on the line attractor.

We next characterize how quickly the evoked activity relaxes back to baseline after the perturbation. Returning to the expression of the perturbation evolution in Fourier space, the propagator can be written in terms of the stability matrix 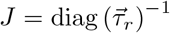 (W−I) (see **Appendix G 4**):

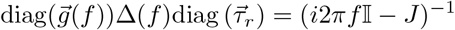

Here the matrix on the left has the same eigenvectors as *J*, meaning they project perturbations in filtered activity space along the same directions though with different magnitudes (e.g. different eigenvalues). If the eigenvalues of −diag(*τ*_*E*_, *τ*_*I*_)^*−*1^Δ(0)^*−*1^ are *χ*(0), then we have the decay time (See **Appendix G 5**):

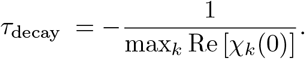

When *τ*_*E*_ = *τ*_*I*_ and the eigenvalues are real, this expression is even simpler as the eigenvalue of the propagator is directly proportional to the relaxation time: 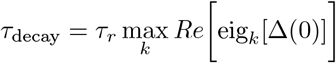. We have that

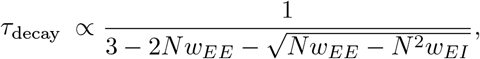

giving *τ*_decay_ explicitly in terms of (*w*_*EE*_, *w*_*EI*_), which produces a monotonically increasing relaxation time along the line attractor (**Fig. 5e** black). This means that networks with pairings high on the line exhibit longer lasting perturbations than those with pairings low on the line.

To compare this to simulation, we assume 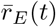 will decay according to an exponential 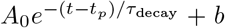 where *A*_0_ is the response amplitude at perturbation time *t* = *t*_*p*_. We quantify a relaxation timescale *τ*_decay_, such that 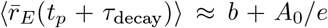. This roughly corresponds to the amplitude of the decay after a full time constant has elapsed. We see the simulation relaxation time (**Fig. 5e** grey) is in good agreement with the mean-field model.

#### Pattern Completion

Additionally, recurrently coupled networks are known to exhibit pattern completion [95–97], which measures the propensity of a network to respond when only a subset of the neurons are stimulated. We test this on the networks that are high or low along the line attractor by stimulating 20% of the excitatory neurons. Unsurprisingly, we find that the network higher on the line attractor exhibits a stronger response and thus recruits more unstimulated neurons in the network to respond relative to networks low on the line attractor (**Fig. 5f**).

#### Co-variability and fluctuations

Finally, the propagator gives insight into the magnitude and temporal structure of the co-fluctuations of neural activity across the population. The autocorrelation function of the simulated *E* population activity 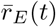 displays a clear progression along the synaptic weight line attractor: its amplitude increases, and its excursions at nonzero lags become larger when moving up the line attractor (**Fig. 5g**; purple to orange traces). Using the covariance of firing rates for the Hawkes process (see **Appendix G 7**), we derive the mean-field cross spectrum (the Fourier transform of the autocorrelation function) (see **Appendix G 8**):

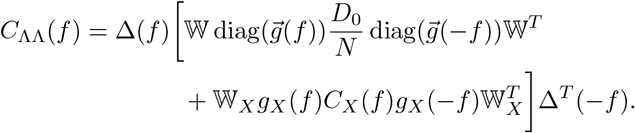

Here 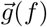 is the Fourier transform of the exponential synaptic kernel filter in the Hawkes process, *D*_0_ is the diagonal 2 2 matrix of the firing rates, and *C*_*X*_(*f*) is the Fourier transform of the mean-field cross-correlation for the external population, which is 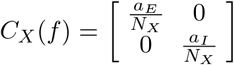.

The inverse Fourier transform of the *EE* component of *C*_ΛΛ_(*f*) gives a mean-field theory estimate of the autocorrelation function of *r*_*E*_(*t*). It shows similar dependency of its shape on the position along the weight line attractor as the Hawkes process network (compare **Fig. 5g** and **Fig. 5h**). Crucially, this dependency is through the propagator, because the input correlation structure (*C*_*X*_(*f*)) and the recurrent firing rates (*D*_0_) do not change along the line, while the propagator Δ(*f*) does.

Finally, we can quantify the trial-to-trial variability of the population-averaged firing rates along the line attractor, as in the previous section (**Fig. 2f**). Mathematically, this corresponds to the inverse Fourier transform of the cross-spectrum evaluated at zero lag, 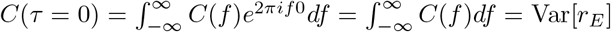. Upon moving up the line attractor, Δ(*f*) becomes large until the rate dynamics become unstable. Thus trial-to-trial variability grows when moving further up the line attractor (**Fig. 5i**).

#### Signal to Noise Ratio

The preceding analysis showed that the propagator amplifies both evoked responses (i.e. signal) and the ongoing co-fluctuations of neuronal activity (i.e. noise). Since both signal and noise are governed by the same operator Δ(*f*), a natural question is which grows faster along the line attractor. That is, as responses become larger along the line attractor, do they become more reliable?

To investigate we compute the signal-to-noise ratio (SNR):

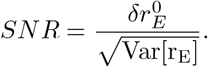

Here 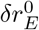 is the peak amplitude response to the perturbation. The SNR is non-monotonic along the line, reaching a peak a little before halfway up the line (**Fig. 5j**). Intuitively, we can understand why this occurs. The variance increases supra-linearly (**Fig. 5i**), while the gain increases linearly (**Appendix G 2**). Initially, Var [*r*_*E*_] increases very slowly along the line, allowing initial response amplitude (signal) to dominate. Eventually though, the rapid growth of variability dominates as the propagator approaches its singularity.

Altogether, we see that the position along the synaptic line attractor ‘parameterizes’ how strongly and how persistently a given excitatory population reacts to transient perturbations, even though its mean baseline activity is indistinguishable across the line attractor. This suggests a natural way for the same assembly to exist in multiple, graded forms: networks at different points on the line encode the sensitivity or salience of a learned pattern, not just its presence or absence [9, 63].

Along the line, these coupled effects imply a tradeoff: stronger assemblies (stronger and longer response) are accompanied by larger shared fluctuations. The resulting nonmonotonic SNR highlights an intermediate regime where responses, and thus recurrent amplification of network activity, are large yet remain comparatively reliable, before variability diverges near the stability boundary. This suggests there is an intermediate “sweet spot” in which a network response will be optimally coherent relative to the ongoing co-fluctuations. In fact, if all external neurons are perturbed and forced to spike (as opposed to the recurrent neurons in the previous analyses), the SNR decreases monotonically. If the goal is to amplify a signal from a perturbed external source, then stronger connectivity will not make the signal clearer (**Supplement Fig.S2**).

### Effect of Correlations on the line attractor weight dynamics

Networks of spiking neurons can operate in one of two distinct dynamical regimes: the *asynchronous state* or the *correlated state*. The extent to which the cortex is in either of these states is debated, but both are pervasive [69, 98, 99]. So far, we have considered our network to be in the asynchronous state, characterized by the absence of significant correlations between spike trains (**Fig. 1d**).

However, neuronal circuits *in vivo* can also operate in the correlated state [69, 100, 101]. Correlations between spike trains typically either follow from strong recurrent connections [12, 91, 92, 102], intrinsic plasticity via elevated excitability [103, 104], or common external input [69, 74]. Because Hebbian plasticity strengthens synapses between co-active neurons, these spike train correlations can become sufficiently large to further shape wiring [89, 105, 106]. Thus, we next ask how correlations affect the weight dynamics and the existence of a line attractor in weight space.

In order to do this, we consider external synchronous fluctuations as the source of pairwise correlations in our model. We add a shared Ornstein–Uhlenbeck (OU) component *η*_*α*_(*t*) to the intensity functions of the external populations of either Λ_*EX*_ or Λ_*IX*_ (**Fig. 6a**):

**FIG. 6.**
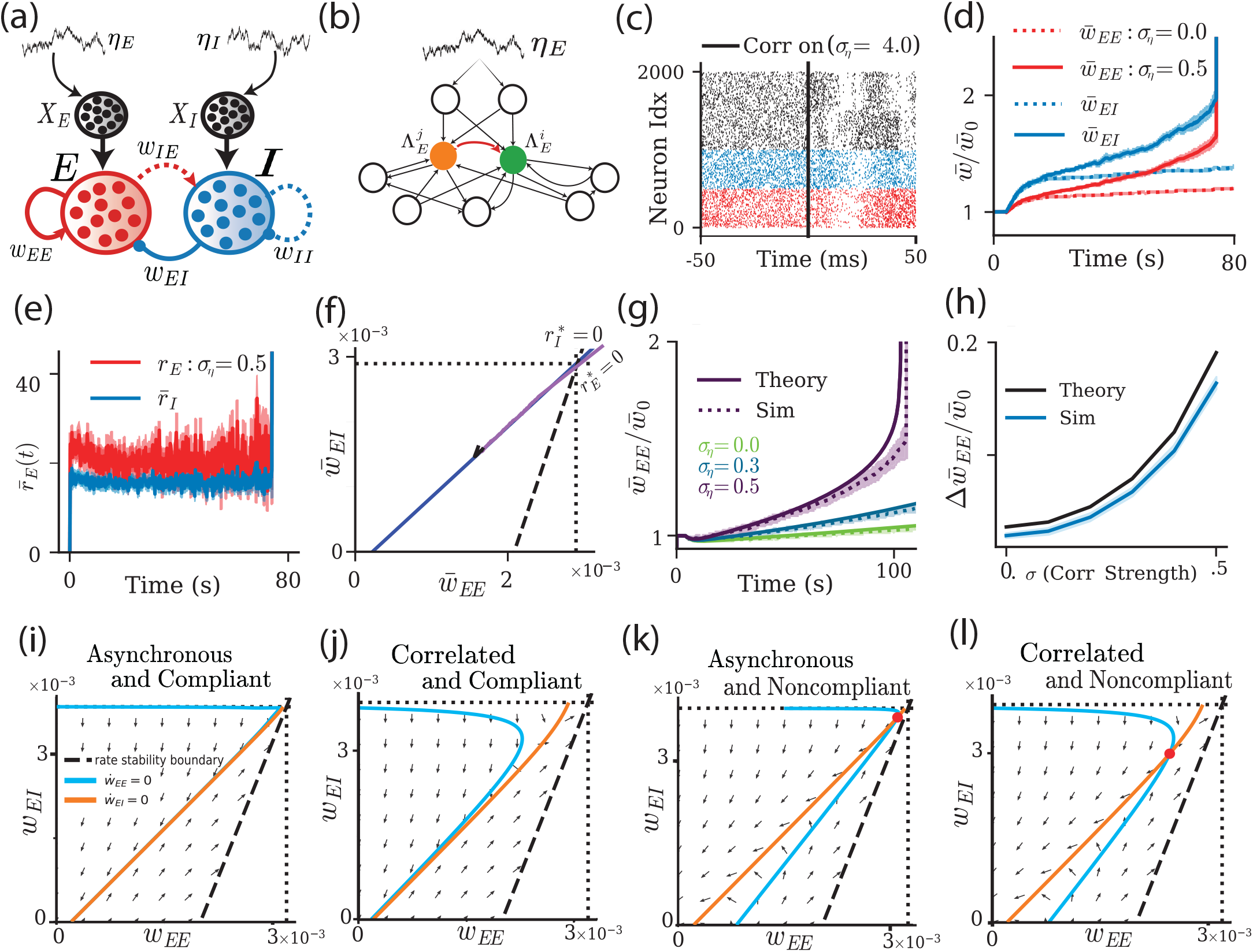
The effect of correlated spiking activity on synaptic weight dynamics. **a**. Schematic of the recurrent *EI* network where neurons in the external populations *X*_*E*_ and *X*_*I*_ receive a shared common fluctuating input (*η*_*E*_ (*t*) and *η*_*I*_ (*t*)). **b**. Two neurons from the external population driving the *E* population with intensity functions 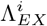 and 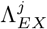 receive common fluctuating input *η*_*α*_(*t*). **c**. Spike train raster plot of *E* (red), *I* (blue), and *X* (black) neurons. Vertical black line marks the time point when correlations are applied (*σ*_*η*_ = 4). **d**. Weight dynamics without (*σ*_*η*_ = 0, dotted lines) and with fluctuating inputs (*σ*_*η*_ = 0.5, solid lines) for *E* (red) and *I* (blue) weights. **e**. Firing rate dynamics for networks with common fluctuating inputs applied to *E* (red) and *I* (blue) rates (*σ*_*η*_ = 0.5). **f**. Weight dynamics in the (*w*_*EE*_, *w*_*EI*_) phase space without (*σ*_*η*_ = 0, black) and with common fluctuating inputs (*σ*_*η*_ = 0.5, purple line). The blue line marks the line attractor when *σ*_*η*_ = 0. **g**. Change of normalized 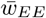 over time for different values of amplitudes *σ*_*η*_. Dotted colored lines indicate the Hawkes process network model, while solid lines indicate the mean-field model. **h**. Normalized weight change 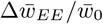 from 25 to 70 s as a function of correlation strength *σ*_*η*_ for the Hawkes process model (blue) and the associated mean-field model (black). Shading in d and e denotes the SEM across 10 trials; in g it denotes the SD across 10 trials, and in h the SEM across 10 trials. **i**. The flow fields and nullclines for the slow system in the asynchronous and compliant state. **j**. Same as i, but for the correlated and compliant state. **k**. Same as i and j, but for the asynchronous and noncompliant state. The red dot indicates a new fixed point in which the nullclines intersect. **l**. Same as i,j, and k, but for the correlated and noncompliant state.

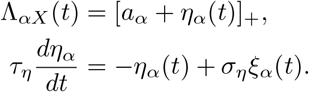

Here *α* ∈ {*E, I*}, *ξ*_*α*_(*t*) is a Gaussian white noise process, *τ*_*η*_ and *σ*_*η*_ are the respective time constant and intensity of the OU process, and []_+_ is a rectifier function. The process *η*_*α*_(*t*) models a temporal inhomogeneity in the firing rates of the individual external neurons, making the external spike trains 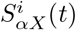 a doubly stochastic process [107, 108]. The fact that *η*_*α*_(*t*) is shared among all external spike trains driving population *α* will cause significant correlations in the external inputs to pairs of neurons (**Fig. 6b**). These increased correlations result in spiking synchrony, which can be seen in the raster plot after adding the OU process (**Fig. 6c**).

Without external correlations (*σ*_*η*_ = 0), the mean synaptic weights, 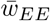 and 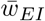 both reach a steady state on the line (**Fig. 6d**; dotted lines). By contrast, when external inputs are correlated (*σ*_*η*_ = 0.5), resulting in significant pairwise *E* spiking correlations, there is a positive deterministic drift in the mean synaptic weights 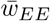 and 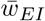 (**Fig. 6d**; solid lines). This suggests that the line attractor dynamics present for *σ*_*η*_ = 0 are no longer in effect. Indeed, this drift eventually drives the network towards instability (**Fig. 6d**; solid lines at *t* ≈ 80s). However, before the network activity becomes unstable the mean firing rates remain fixed because of the homeostatic control through *w*_*EI*_ dynamics (**Fig. 6e**). This suggests that while the weights are drifting in the network with correlated spiking activity, they are drifting along what is the line attractor for the asynchronous network (**Fig. 6f**). We remark that in the previous results in our Hawkes process network model (**Figs. 1, 2**), this (slow) drift was absent because we focused our analysis on intermediate timescales (~30s). Longer timescales, however, reveal that the Hawkes process network is not perfectly asynchronous, but that the recurrent *E* weights induce weak pairwise correlations [79].

To understand this drift in weight space, we extend the mean-field model to accommodate spike train correlations following previously published approaches [79]. The weight dynamics in the mean-field model now depend on additional terms (see **Appendix B** and **G 8** for details):

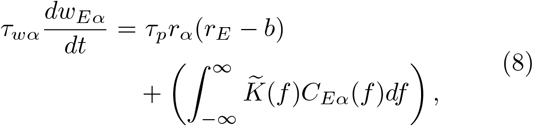

where *α* ∈ {*E, I*}. Here 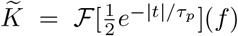 is the Fourier transform of the STDP kernel, which is integrated against the spike train cross spectrum of the recurrent units *C*_*Eα*_(*f*). This term scales as O(*N* ^*−*1^) in the asynchronous regime, and can thus be neglected in the large *N* limit (see Eq. 3). However, in the correlated regime the term does not vanish in the large *N* limit and must be considered. The mean-field spike-count crossspectrum of the network is:

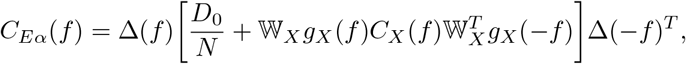

with 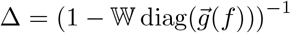 being the propagator as defined in the past section, and *D*_0_ is the diagonal matrix of the mean firing rates. Note, *C*_*Eα*_ refers to the spikecount cross spectrum, whereas in the previous section, we used the intensity function cross spectrum *C*_ΛΛ_ for the trial-to-trial variability. The former conveys the spiketrain correlations, while the latter conveys the correlations of the underlying firing rates. The function *C*_*X*_(*f*) is the Fourier transform of the mean-field cross correlation for the external population, defined as:

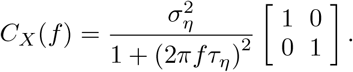

We note that *C*_*X*_(*f*) = 0 if *σ*_*η*_ = 0.

Correlations induce a drift in (*w*_*EE*_, *w*_*EI*_) space through the integral term in Eq. (8). This drift destroys the synaptic weight line attractor observed in the asynchronous state. This is because the first term in Eq. (8) dominates the initial dynamics, and upon reaching the line attractor defined for *σ*_*η*_ = 0 (where *r*_*E*_ ≈ *b*), the contribution of the first term is close to zero. Thus, subsequent dynamics are due to the correlation term, resulting in a deterministic drift, while firing rate homeostasis is maintained. The drift is positive because the integral of the STDP kernel is positive and the pairwise correlations are positive. We plot the mean synaptic *w*_*EE*_ weight for different noise amplitude (*σ*_*η*_) values, where this drift becomes faster as *σ*_*η*_ increases for both the mean-field model (**Fig. 6g**; solid lines) and the Hawkes process network model (**Fig. 6g**; dotted lines). Finally, we calculate the Δ*w*_*EE*_ over the interval from 25 to 70 s, where *w*_*EE*_ increases linearly. We see the drift grows supra-linearly with *σ*_*η*_, matching well between simulations of the Hawkes process and associated mean-field theory (**Fig. 6h**).

To understand this drift and sources of instability, we first consider the *w*_*EE*_ and *w*_*EI*_ nullclines for the network with asynchronous spike trains and where the *E*→ *E* plasticity is homeostatically compliant (blue and orange curves, **Fig.6i**). In this case, the *w*_*EE*_ nullcline is the union of the 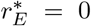 and 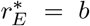 curves in synaptic weight space. Further, the *w*_*EE*_ and *w*_*EI*_ nullclines overlap, meaning that a line attractor is present. In contrast, when the network spiking activity becomes correlated, the two *w*_*EE*_ nullcline curves merge into one nonlinear curve, which no longer overlaps with the *w*_*EI*_ nullcline (blue curve, **Fig. 6j**; *σ*_*η*_ = 1.5 is used to show this clearly). This removes the line attractor and leads to dynamical flow up and to the right in (*w*_*EE*_, *w*_*EI*_) space.

We next consider the necessity of homeostatic compliance for line attractor dynamics. For both the networks with asynchronous and correlated spike trains where the *E* → *E* synapses are not homeostatically compliant, the *w*_*EE*_ and *w*_*EI*_ nullclines intersect at a saddle point (**Fig. 6k, l**). Consistent with **Fig. S1d**, this would lead to trajectories decaying or blowing up depending on which side of the nullclines an initial condition is set. In total, line attractor dynamics require both asynchronous spike trains and homeostatically compliant *E* → E plasticity.

In summary, inducing spike train correlations in the recurrent network via external shared fluctuations to each population causes drift in the synaptic weights along where the line attractor would exist if correlations were absent. Accounting for pairwise spike time correlations in our mean-field model can capture the dynamics of the deterministic drift in synaptic weights. If this drift is left unchecked, it will lead to a blowup of *E* weights and rates. This seriously undermines the ability of the network to store information and maintain persistent population activity. We thus conclude that the network is both sensitive to potential competition between plasticity rules with different plasticity thresholds and to correlations driving the system to runaway excitation. In the final section, we will present one possible solution to overcome this unbounded drift in the presence of pairwise correlations.

### Correlations between E and I cells can mitigate synaptic weight Drift

In cortical circuits, it is well established that both excitatory and inhibitory neuronal populations often receive common inputs, such as shared thalamic or intracortical drive in sensory cortices [109, 110]. This has been shown to drive the system back towards asynchrony [99]. We induce correlations across *E* and *I* neurons by correlating their external populations via their intensity functions:

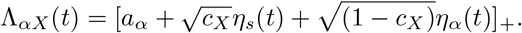

Here, *η*_*s*_(*t*) and *η*_*α*_(*t*) are OU processes, and *c*_*X*_ is the correlation coefficient between these external inputs, which sets the proportion of the noise that is shared or private between the *E* and *I* populations (**Fig. 7a,b**). For simplicity, they have the same noise intensity *σ*_*η*_. This results in a new external correlation structure in the meanfield model of the form:

**FIG. 7.**
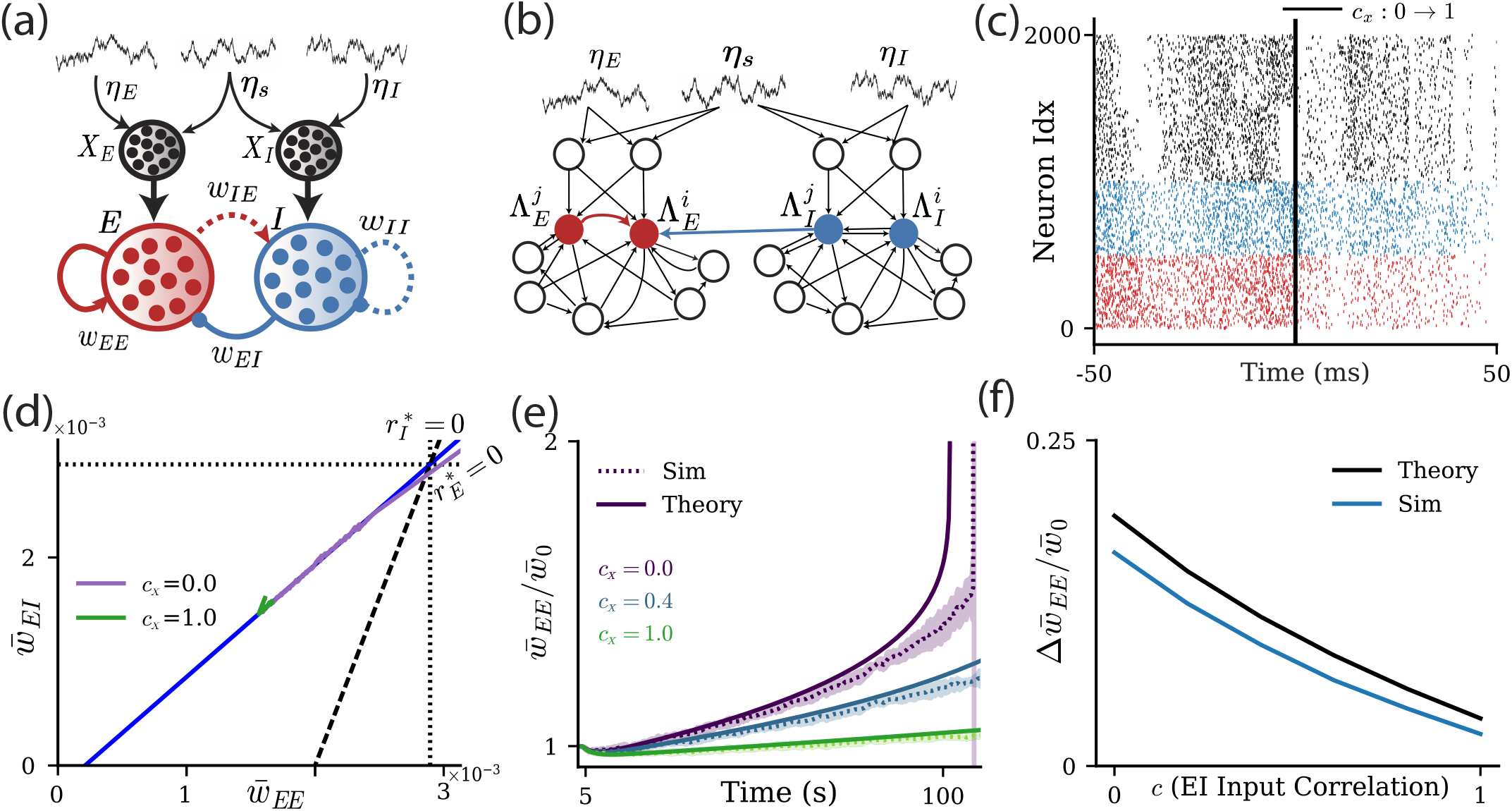
Correlations to both *E* and *I* populations can mitigate synaptic weight drift. **a**. Schematic of the network when the external inputs to *E* and *I* populations are now correlated. The parameter *c*_*X*_ denotes the proportion of input shared between the external populations which activate *E* and *I* together. **b**. Schematic of the ensemble in which the fluctuating inputs can now be shared between the external populations which synapse onto *E* and *I. I → E* connections (blue) and *E → E* connections (red) between example pairs of neurons can potentiate or depress. **c**. Spiking raster of *E* (red), *I* (blue), and *X* (black) neurons. Vertical black line indicates when correlations between *E* and *I* with *c*_*X*_ = 1 are activated. **d**. Traces of the mean synaptic weights in (*w*_*EE*_, *w*_*EI*_) phase space showing drift towards the instability when correlated inputs to *E* and *I* are purely private (*c*_*X*_ = 0, purple curve) vs shared (*c*_*X*_ = 1, green curve). **e**. As *c*_*X*_ increases from 0 to 1, the drift is mitigated, so the mean synaptic weight drifts less and less. **f**. Drift for simulations of the Hawkes process network (blue) and associated mean-field theory (black).

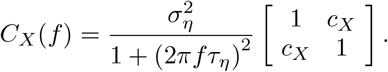

Adding shared fluctuations to the *E* and *I* populations leads to a reduction of pairwise *E* → *E* spike time correlations (see spike train raster in **Fig.7c**). The drift in mean weight dynamics observed with only private correlations (**Fig. 7d**; purple curve) is suppressed when the correlations are shared between *E* and *I* populations (**Fig. 7d**; green curve). For both simulations of the Hawkes process network (**Fig. 7e**, dotted curves) and associated meanfield theory (**Fig. 7e**, solid curves), we see the slope of the 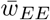 evolution gets smaller as *c*_*X*_ approaches one (**Fig. 7e**). Indeed, when *c*_*X*_ = 1 the drift is similar to that seen in the case of *σ*_*η*_ = 0 (when pairwise correlations are negligible). Finally, we find the drift rate monotonically decreases as *c*_*X*_ varies from 0 to 1 for both theory (**Fig.7f**, black curve) and Hawkes process simulations (**Fig.7f**, blue curves).

In summary, by sharing the fluctuations between the *E* and *I* populations, we induce a cancellation of the external correlations projecting onto the *E* population by the recurrent *I* population fluctuations onto the *E* population. As a result, for large *c*_*X*_ the spiking activity of the *E* and *I* populations is approximately asynchronous. Because of this the stability of weight dynamics is similar to the case when external inputs are uncorrelated (*σ*_*η*_ = 0).

## III. DISCUSSION

A central question in assembly learning is: how can recurrent circuits turn transient correlated activity into a stable, strongly coupled circuit? In the classic cell-assembly picture, co-active excitatory neurons strengthen their recurrent connections, making future coactivation more likely and thereby allowing a stimulus, memory, or task variable to be encoded. However, the same positive feedback that makes assemblies responsive to inputs also makes them potentially unstable: recurrent excitation can amplify activity, correlations, and future synaptic potentiation. Thus, proper assembly learning cannot consist simply of forming strong *E* → *E* connections, but must involve maintaining a stable operating point and preserving useful dynamical responses.

Existing assembly models often conceptualize learning as a transition from an unstructured network to a fully formed assembly. While this dichotomy is useful, it leaves open a computational question: once an assembly exists, how might its connectivity shape circuit function? In our study we expanded on past work [37, 53] and explored how the interaction of *I* → *E* homeostatic plasticity and *E*→ *E homeostatically compliant* plasticity can not only stabilize assembly learning, but also allow for flexibility of assembly learning along a continuum.

More broadly, homeostatic compliance is necessary when multiple plasticity mechanisms act on the same postsynaptic neuron; their fixed points and timescales cannot be considered independently. Agnes and Vogels [38] showed that conventional excitatory and inhibitory rules with mismatched firing-rate set points can compete, causing synaptic weights to diverge or vanish. By contrast, co-dependent rules relax this constraint by making inhibitory plasticity depend on local excitatory (NMDA-mediated *Ca*^2+^) and inhibitory (GABA-mediated chloride) currents. Inhibition then dynamically tracks excitation to maintain a target *E/I* ratio, avoiding the need for matched firing-rate fixed points.

The condition for homeostatic compliance, *b*_*EE*_ = *b*_*EI*_, represents an idealized scenario whose biological basis has not been established. Rate-dependent homeostatic rules are often described by a formalism introduced by Bienenstock, Cooper and Munro (BCM) [26]. In the original BCM theory, the LTP/LTD threshold was a slowly adapting function of recent postsynaptic activity, and subsequent modelling work has proposed mechanisms involving NMDA-receptor-dependent calcium signaling, metaplasticity, or intrinsic excitability [26, 87]. Consistent with this possibility, experimental work found that pairing-induced plasticity at both excitatory and inhibitory synapses onto layer 5 auditory-cortical pyramidal neurons depended on NMDA receptors and L-type *Ca*^2+^ channels [40]. This shared dependence suggests that a common postsynaptic calcium signal could coordinate the two plasticity thresholds while retaining synapse-specific downstream expression. Thus, *b*_*EE*_ = *b*_*EI*_ could plausibly arise if they are determined by a common slowly varying postsynaptic activitydependent signal, such as intracellular Ca^2+^. However, current evidence does not establish that these thresholds are universally coupled.

Further, the exact homeostatic compliance condition of *b*_*EE*_ = *b*_*EI*_ is unlikely to occur in nature, and an approximate condition of *b*_*EE*_ ≈ *b*_*EI*_ will not result in a line attractor in synaptic weight dynamics. Nevertheless, *b*_*EE*_ ≈ *b*_*EI*_ can lead to slow passage effects as synaptic weights converge to an isolated fixed point, resulting in dynamics that transiently show a continuum of synaptic weights. However, the nullcline structure for *b*_*EE*_ ≠ *b*_*EI*_ produces a fixed point at high (*w*_*EE*_, *w*_*EI*_) pairings (**Fig. 6k**), meaning that slow passage effects will be more apparent for strong synaptic weights compared to weak weights.

A consequence of using *I* → *E* plasticity as a homeostatic mechanism is the continuum of stable network configurations organized by a line attractor in synaptic weight space. While previous rate-based models characterized how interacting excitatory and inhibitory plasticity can generate line attractors in synaptic weight space [37, 53], those studies did not address what functional role such line attractors could enact. Related work on homeostatic structural plasticity showed that engram strength can be encoded by position along a connectivity line attractor in excitatory weight space and can monotonically increase pattern completion [111]. A key finding in our study is that different synaptic pairings along the line attractor correspond to distinct dynamical response properties despite having the same steady state firing rate. Along the attractor, the network changes how it amplifies inputs, filters fluctuations, and responds across timescales. In this sense, the line attractor does not store persistent activity, as in classical working-memory models [112, 113], but instead stores a continuum of synaptic configurations that shape how the circuit processes timevarying inputs.

Previous assembly models have associated strengthening of recurrent assembly structure with improved performance on particular coding and recall measures [13, 111]. Our results give an alternate viewpoint. We found that the strongest assembly is not necessarily the most computationally useful: greater recurrence increases gain, but it also amplifies shared fluctuations. This leads to a degradation in the SNR for strongly coupled assemblies. Reliable transmission may therefore be optimized at intermediate assembly strength in our model, rather than maximal recurrence.

This continuum of network configurations which shape signal processing also provides an interpretation of *learning* within our framework. Fluctuations in fast firingrate dynamics drive slower synaptic changes, producing a gradual drift along the former line attractor toward a stronger assembly structure. In our model, such learning is associated with transient violations of *E/I* balance, consistent with experimental work suggesting that disruptions of *E/I* balance can facilitate *E* synaptic plasticity [44, 45, 114].

At the same time, persistent drift can eventually drive the network to instability. Because recurrent networks are never perfectly asynchronous, weak correlations can slowly drive synaptic potentiation even in the absence of structured input [13, 79, 89]. Additional correlations from shared external excitatory input further accelerate this process. Our results, therefore, suggest that mechanisms promoting asynchronous activity and decorrelation may play an important role in stabilizing long-term assembly dynamics. We suggest correlations between *E* and *I* populations as a mechanism to mitigate such drift, as seen experimentally [99, 110]. Possible alternative candidate mechanisms to ensure asynchronous network activity include inhibitory feedback, sparse connectivity, and short-term synaptic depression [70, 92, 115, 116].

There can be many sources of drift along the line attractor, with the direction determined by the average rate- and correlation-dependent contributions to synaptic plasticity. In our model, positive *E* → *E* correlations combined with a potentiation-dominated STDP kernel led to a positive drift. However, a depression-dominated plasticity or anti-correlations between distinct excitatory neurons could both lead to a negative drift down the line. Intuitively, this would be due to the drift term 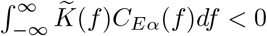, leading to a negative drift. Adionally, just as excitatory pulses induce coordinated potentiation (Fig. 2), inhibitory pulses can induce coordinated depression. Thus, environmental fluctuations have no universal preferred direction. Instead, the net direction depends on how correlations interact with the balance of potentiation and depression.

We interpret our *EI* network as a single candidate assembly, whose neurons have already been allocated, embedded within a larger recurrent circuit. This starting point is biologically plausible because elevated intrinsic excitability or CREB activity can bias neurons into a memory trace, after which recurrent connections among the recruited neurons can be potentiated [117, 118]. However, both the surrounding circuit (*X*) and the recurrent *E* → *I* and *I* → *I* pathways are simplified to be nonplastic. This abstraction permits a mean-field model with only 2 dynamical variables (*w*_*EE*_, *w*_*EI*_) vs. 8+, which would be the case if multiple assemblies with fully plastic connections were to be considered. Extending the framework to include simultaneous plasticity within and between multiple allocated populations will be an important direction for future work. In particular, it would be interesting to investigate how multiple line attractors in weight space could be combined to implement more complex computations, for example, in the setting of a ring attractor [119]. More broadly, the distinct response properties that emerge along the line attractor may provide a useful substrate for coordinating assemblies across multi-area networks to support task-relevant computations [120, 121].

Finally, although our framework permits ongoing synaptic plasticity, it assumes fixed baseline firing rates. Previous work has explored the emergence of heterogeneous firing rates in networks with homeostatic inhibitory synaptic plasticity [38, 122]. Extending our framework to allow heterogeneity in both firing rates and synaptic weights could therefore provide further insight into the computational advantages of flexible assembly organization.

## ACKNOWLEDGMENTS

B.D. is supported by the NIH grants R01EY037119, R01NS133598, T90DA059109; NSF grant 2605555, and the Simons Foundation Collaboration on the Global Brain.

C.M. was funded by a Human Frontier Science Program Postdoctoral Fellowship LT0005/2024 L (CM).

This research benefited from the Physics Frontier Center for Living Systems funded by the National Science Foundation (PHY-2317138). Financial support was provided via the National Institute for Theory and Mathematics in Biology (Simons Foundation award MP-TMPS-00005320 and NSF award DMS-2235451).

## IV. METHODS

### A. Model parameters

#### Hawkes process network model

Using a framework developed by Hawkes of interacting self-exciting point processes [71, 72], we construct a model of two populations of spiking neuron models with *N*_*E*_ excitatory and *N*_*I*_ inhibitory neurons, which are connected within and across populations. The intensity functions for the *E* and *I* populations, i.e. the instantaneous rate at which events occur from a Poisson point process, are modeled as:

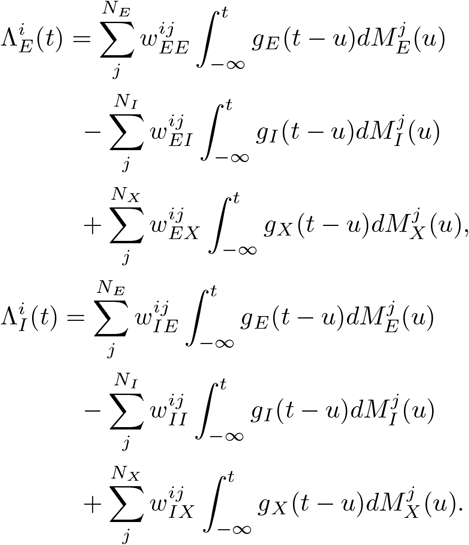

**TABLE I.**
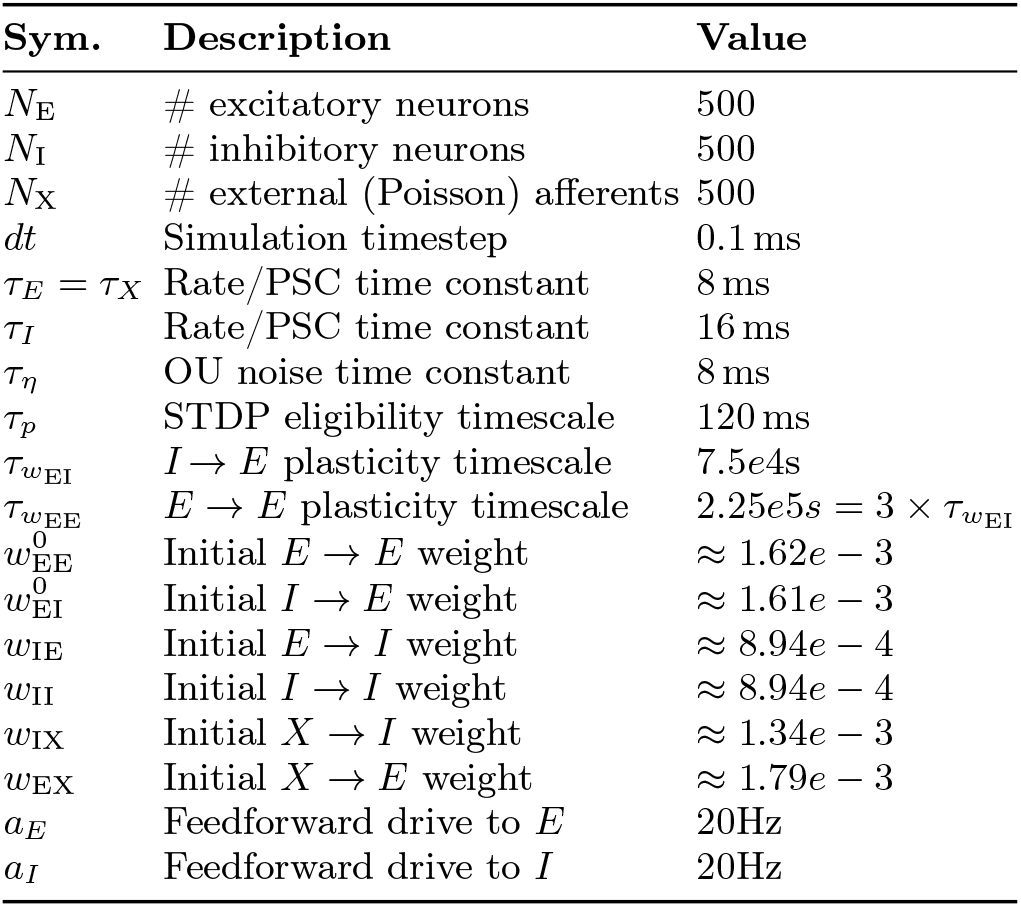
Parameters used across all figures for the Hawkes process network and associated mean-field models (unless otherwise specified).

| **Sym.** | **Description** | **Value** |
|---|---|---|
| *N*<sub>E</sub> | # excitatory neurons | 500 |
| *N*<sub>I</sub> | # inhibitory neurons | 500 |
| *N*<sub>X</sub> | # external (Poisson) afferents | 500 |
| *dt* | Simulation timestep | 0.1 ms |
| τ<sub>*E*</sub> = τ<sub>*X*</sub> | Rate/PSC time constant | 8 ms |
| τ<sub>*I*</sub> | Rate/PSC time constant | 16 ms |
| τ<sub>η</sub> | OU noise time constant | 8 ms |
| τ<sub>*p*</sub> | STDP eligibility timescale | 120 ms |
| τ<sub>*w*<sub>EI</sub></sub> | *I* → *E* plasticity timescale | 7.5*e*4s |
| τ<sub>*w*<sub>EE</sub></sub> | *E* → *E* plasticity timescale | 2.25*e*5*s* = 3 × τ<sub>*w*<sub>EI</sub></sub> |
| *w*<sup>0</sup><sub>EE</sub> | Initial *E* → *E* weight | ≈ 1.62*e* − 3 |
| *w*<sup>0</sup><sub>EI</sub> | Initial *I* → *E* weight | ≈ 1.61*e* − 3 |
| *w*<sub>IE</sub> | Initial *E* → *I* weight | ≈ 8.94*e* − 4 |
| *w*<sub>II</sub> | Initial *I* → *I* weight | ≈ 8.94*e* − 4 |
| *w*<sub>IX</sub> | Initial *X* → *I* weight | ≈ 1.34*e* − 3 |
| *w*<sub>EX</sub> | Initial *X* → *E* weight | ≈ 1.79*e* − 3 |
| *a*<sub>*E*</sub> | Feedforward drive to *E* | 20Hz |
| *a*<sub>*I*</sub> | Feedforward drive to *I* | 20Hz |

**TABLE II.** Parameter choices for Figure 4.

|  |  |  |
|---|---|---|
| *w*<sub>IE</sub> | Initial *E* → *I* weight in Fig 4d. | ≈ 1.97*e* − 3 |
| *w*<sub>IE</sub> | Initial *E* → *I* weight in Fig 4e. | ≈ 8.94*e* − 4 |
| *w*<sub>IE</sub> | Initial *E* → *I* weight in Fig 4f. | ≈ 1.79*e* − 4 |

To better streamline our mean-field derivation we allow for autaptic connections 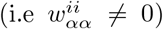; in our meanfield derivation this is equivalent to letting *N*_*α*_ − 1 ≈ *N*_*α*_ (*α ∈* {*E, I*}).

The coefficient 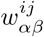 is the coupling strength from neuron *j* in population *β* to neuron *i* in population *α*. 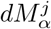 is the differential spike count process for neuron *j* in population *α*. The kernel 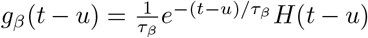, with *β ∈* {*E, I, X*}, integrates over the past spiking activity. The filter time constant is determined by the presynaptic population, with *τ*_*E*_ = *τ*_*X*_ and *τ*_*I*_ = 2*τ*_*E*_. The displayed intensity equations apply when rectification is inactive. The probability of neuron *i* in population *α* producing a spike is approximated as 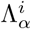 *dt* (for sufficiently small *dt*). In practice, we sample a random number from *U*(0, 1), and if it is smaller than 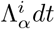 a spike occurs.

The external unit *j* has the intensity function:

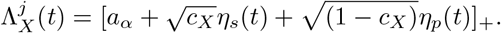

Here, *η*_*s*_(*t*) and *η*_*p*_(*t*) are OU processes, which are shared and private between the external units, respectively. The private OU processes are private between the E- and I-driving populations, but shared within each population. The *η*_*m*_ processes (*m ∈* {*s, p*}) evolve according to 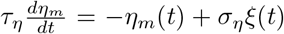, where *ξ*(*t*) is Gaussian white noise, *τ*_*η*_ is the time constant for the OU process, and *σ*_*η*_ scales the size of OU noise (*m ∈* {*s, p*}). The coefficient *c*_*X*_ is the correlation coefficient between external units, which defines the proportion of the noise that is shared between units. We choose a regime in which Λ_*αX*_ ≥ 0 for *α ∈* {*E, I*}.

#### Synaptic plasticity model

Our plasticity model requires eligibility traces of spiking activity: for neuron *j* in population *α ∈* {*E, I*} the trace *x*^*j*^ (*t*) obeys:

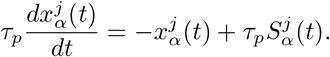

Here, 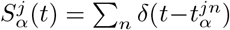 is the spike train from neuron *j* in population *α*. The time constant *τ*_*p*_ sets the temporal selectivity of the spike-timing-dependent plasticity (STDP) process.

Solving the above differential equation (with 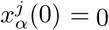 and 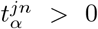 for all *n*) yields 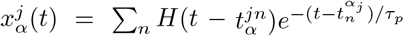. The synaptic weight 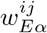 between a presynaptic neuron *j* in population *α* and a postsynaptic neuron *i* in population *E* evolves according to a spiketiming dependent plasticity rule:

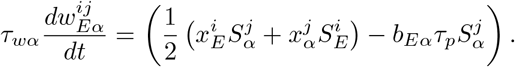

The dynamics of 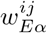 are driven, in part, by combinations of eligibility *x*(*t*) and spiking activity *S*(*t*). The coefficient 1*/*2 reflects the positive symmetric nature of the STDP window (e.g. each half of the STDP window is net LTP). The timescale of weight dynamics is set by *τ*_*wα*_.

#### Linear-response propagator

To compute response properties of networks at different locations along the line attractor, we used the linear response theory of the reduced mean-field rate model (since the Hawkes process model is a linear system then this analysis is not a perturbative one). For these calculations, synaptic weights were held fixed because the rate dynamics evolve on a much faster timescale than the synaptic plasticity dynamics. For a fixed pair of synaptic weights (*w*_*EE*_, *w*_*EI*_), the mean-field rate dynamics can be written in convolution form as:

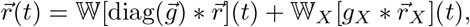

where 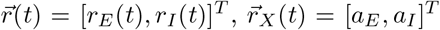, and 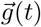 denotes the population-dependent synaptic filters.

In Fourier space, we define the frequency-dependent propagator

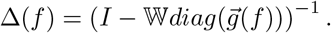

The propagator describes how activity is filtered and recurrently amplified by the network. The equilibrium firing rates are obtained from the zero frequency component,

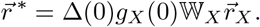

For weights on the line attractor, the excitatory fixed point satisfies 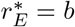.

The same propagator determines the stability of the rate dynamics. For the exponential synaptic filters, the rate dynamics are stable when all eigenvalues of

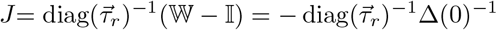

have negative real parts. Here *J* acts on the filtered perturbations 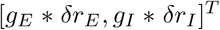, rather than directly on 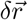. Thus, all response and covariance calculations below were restricted to weight values for which the fixed point 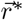 is stable. Equivalently, instability occurs when

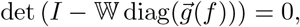

or when the spectral radius of diag 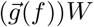*W* reaches one.

### Perturbation protocol

For each location on the line attractor (*w*_*EE*_, *w*_*EI*_) we computed the corresponding steady-state firing rates 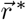. We then injected a brief perturbation into the *E* population and measured the activity deviation from baseline,

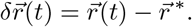

The perturbation was represented as:

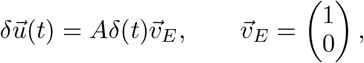

where *A* sets the perturbation amplitude and 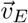 indicates that the input is delivered to the excitatory population. The perturbation represents additional spikes emitted by the E population before synaptic filtering. The response 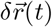 describes the induced change in intensity and excludes the imposed spike pulse itself. The Fourier-domain response is:

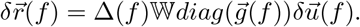

In the time domain, this can be written in terms of the propagator:

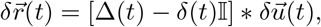

where Δ(*t*) = *δ*(*t*)I + *H*(*t*)W*e*^*Jt*^ diag 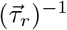. This is the form we use for the theory.

In simulations of the Hawkes process network, the corresponding perturbation was implemented by briefly forcing the stimulated *E* neurons to spike. The population response was then measured from the baseline-subtracted population-averaged *E* firing rate,

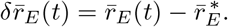

#### Dynamical response metrics

For each (*w*_*EE*_, *w*_*EI*_) point on the line attractor, all response quantities were computed with synaptic weights held fixed. The perturbation response was obtained from the linear-response expression

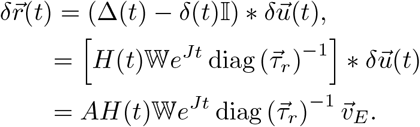

Here, 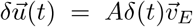. The excitatory component *δr*_*E*_(*t*) was used to compute the time-integrated squared response

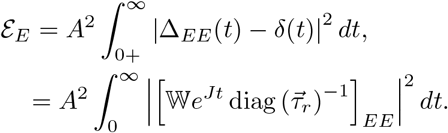

The lower integration limit excludes the instantaneous delta-function component of the propagator and includes only the causal response following the perturbation. We remark that in the time domain the propagator becomes: Δ(*t*) = *δ*(*t*)I + *H*(*t*)W*e*^*Jt*^ diag 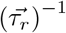, necessitating the −*δ*(*t*)I term in the first line.

The relaxation time of the perturbation was computed from the eigenvalues of the linear response operator at zero frequency. Specifically, if *χ*_*k*_(0) are the eigenvalues of – diag 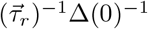, then

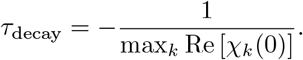

This quantity was evaluated for each (*w*_*EE*_, *w*_*EI*_) pair on the line attractor.

#### Response variability

The covariance and trial-to-trial variability were computed from the mean-field cross spectrum of the population intensity. For the covariance and variance calculations, the spectrum was evaluated as:

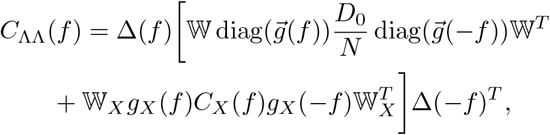

The mean-field autocovariance of the excitatory population was obtained by taking the inverse Fourier transform of the *EE* component of *C*_ΛΛ_(*f*),

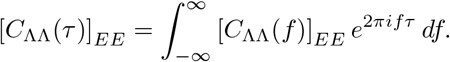

The corresponding trial-to-trial population-rate variance was computed from the zero-lag value (*τ* = 0) of this covariance,

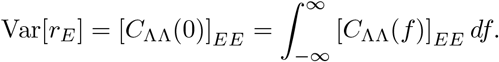

Finally, the signal-to-noise ratio was computed as

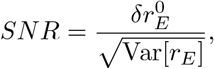

where 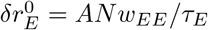 is the peak excitatory response amplitude measured from the same theoretical perturbation response *δr*_*E*_(*t*).

## Code availability

All code used to run the simulations, perform the analyses, and generate the figures presented in this study is publicly available on GitHub at *https://github.com/MikiMiller95/Homeostatic-Tunable-Assemblies*. The repository contains the simulation, analysis, and plotting scripts required to reproduce the results reported here, together with documentation describing the workflow for generating each figure.

## SUPPLEMENTARY FIGURES

**FIG. S1.**
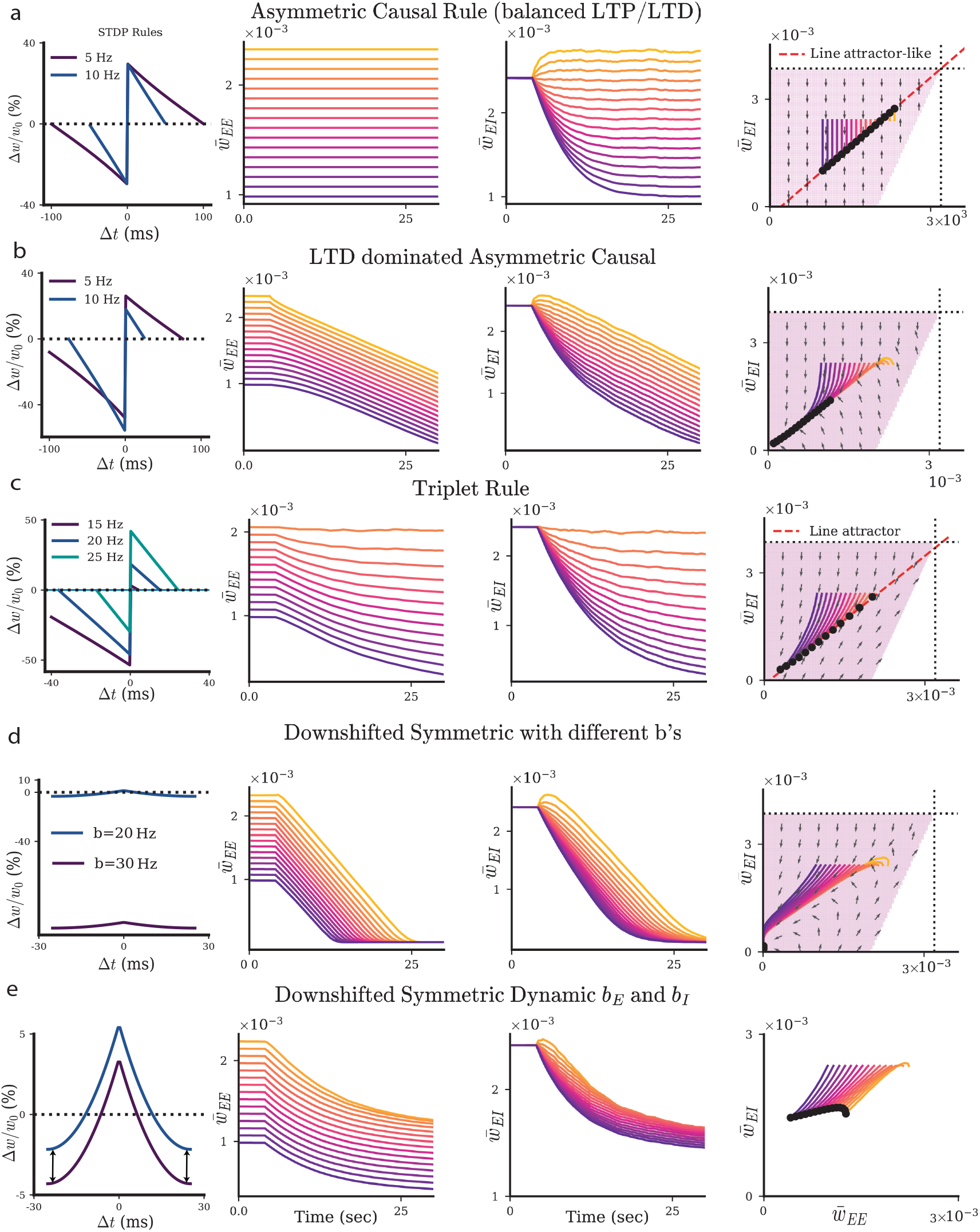
Synaptic Weight Dynamics for Different Learning Rules. **a**. Mean synaptic 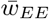 and 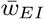 evolution over time for the Asymmetric *E → E* STDP rule (far left; the colors denote different *E* neuron firing rates) with homeostatic *I → E* STDP. In 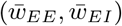 space, they lie on the same line, but 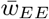 remains constant 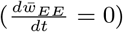. **b**. Mean synaptic 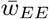 (left) and 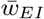 evolution over time for the Asymmetric *E → E* STDP rule but LTD is larger than LTP. The corresponding *E* plasticity rule, 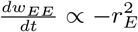, results in persistent depression for 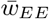 and 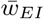. **c**. Mean synaptic 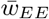 (left) and 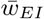 evolution over time for the Asymmetric *E → E* STDP rule which includes triplets of spikes. In 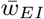 vs 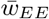 space, they lie on the same line as that found in the pairwise rules used in the text. The corresponding *E* plasticity rule is 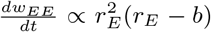. **d**. The plasticity rule in which *b*_*EE*_ = 30 and *b*_*EI*_ = 20 results in net depression for the *E → E* connections, which further drives depression for the *I → E* connections. The mean-field plasticity rule is 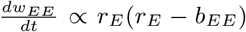. **e**. The *E → E* STDP rule is similarly downshifted initially, but *b*_*EE*_ and *b*_*EI*_ are both allowed to be dynamic according to low-pass filtering of the firing rate 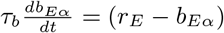.

**FIG. S2.**
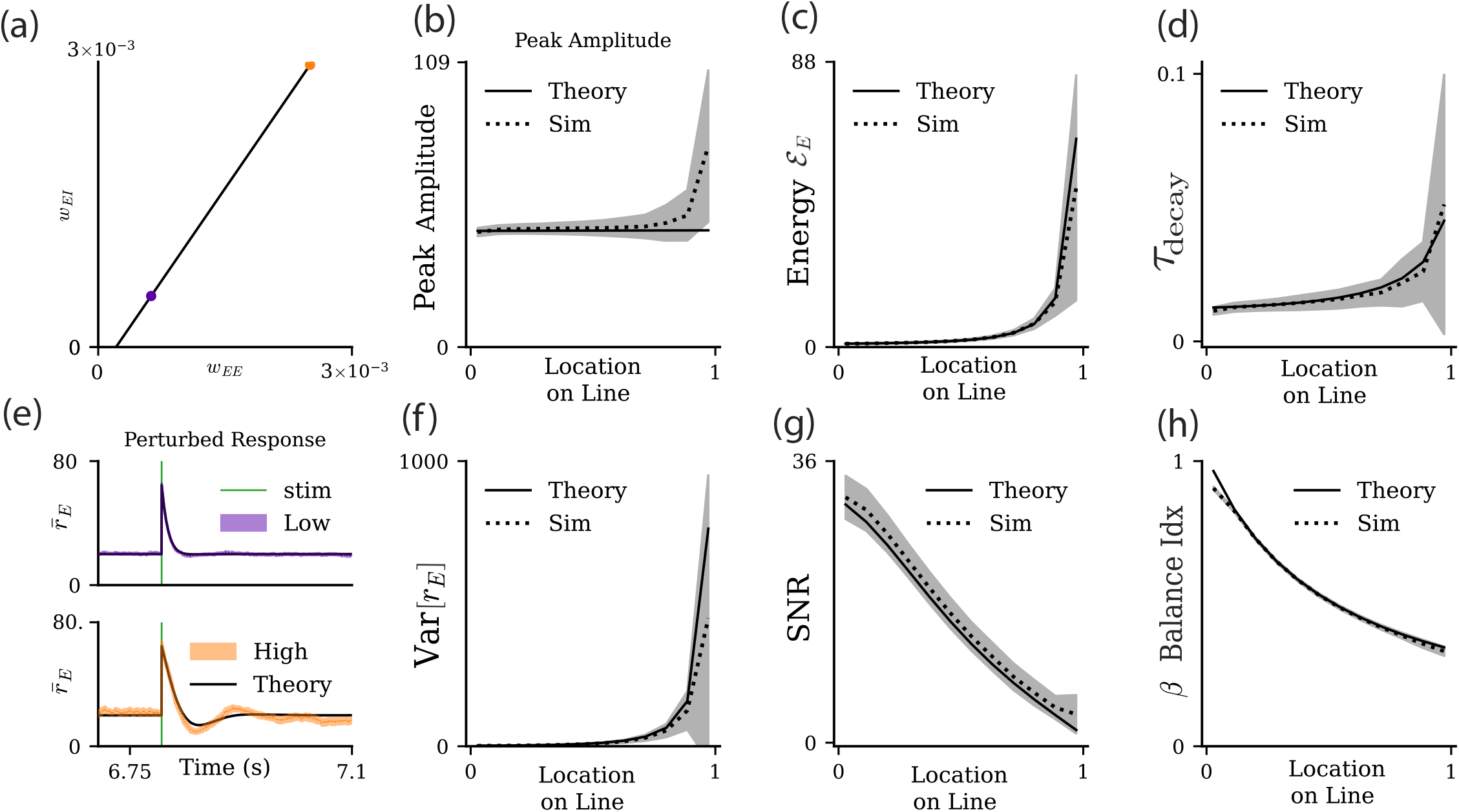
Response properties of the Hawkes process network when the external synapses are perturbed. **a**. Networks at low (purple dot) and high (orange dot) positions on the line attractor are selected. **b**. We similarly parametrize location on the line attractor varying from 0 to 1 starting with where the line attractor crosses *w*_*EE*_ = 0 to the coordinate 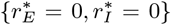. The peak amplitude of the response along the line attractor is flat until it grows nonlinearly near the top of the line attractor. Grey denotes the standard deviation across 20 simulation realizations. **c**. Similar to Figure 5d, the energy ℰ_*E*_ grows along the line attractor for theory (black solid) and simulations (black dashed) along the line attractor. **d**. The relaxation time to a perturbation, *τ*_decay_, grows along the line attractor for theory (black solid) and simulations (black dashed) along the line attractor. **e**. On the line attractor, both networks still fire at the target firing rate *b*. The network with weaker coupling has a short, rapid response (top). Dark purple is theory, light purple denotes simulations. Upon receiving a perturbation, the network with stronger recurrent coupling has a longer rebound in response (bottom). Dark orange is theory. Light orange denotes simulations. **f**. The trial-to-trial variability, Var[*r*_*E*_], grows nonlinearly along the line attractor for theory (black solid) and simulations (black dashed) along the line attractor. **g**. The signal-to-noise ratio now decreases monotonically. The signal of the perturbation in the linear theory is no longer proportional to the connectivity; therefore, stronger connectivity leads to noise dominating more. **h**. The balance index decreases monotonically for both theory and simulations on the line attractor.

## APPENDICES

### A. Derivation of the mean-field rate equations

Before considering the case of the network with plastic weights, we can first consider the mean-field dynamics of this network with fixed weights. Let us consider a typical neuron *i* in population *α ∈* {*E, I*}, whose trial-averaged activity, 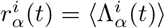, obeys:

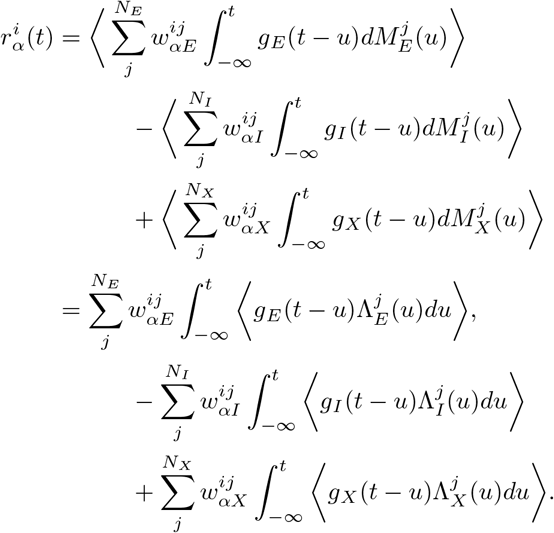

We obtain an evolution equation for 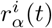 rates by differentiating to yield:

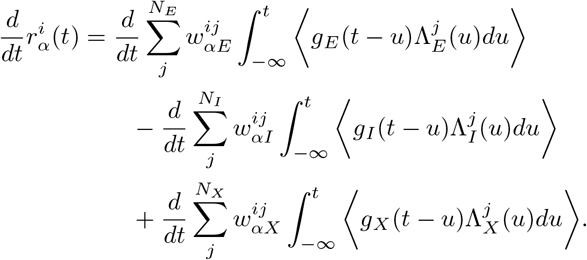

It is noteworthy to acknowledge that the counting process is not actually differentiable; however, we can take a trial average and observe that 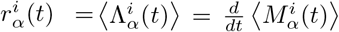. Also consider that 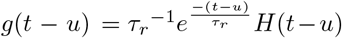 such that 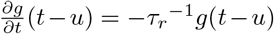. Likewise, we need to consider that we are differentiating a convolution with a variable in the bounds. This will require Leibniz’s rule:

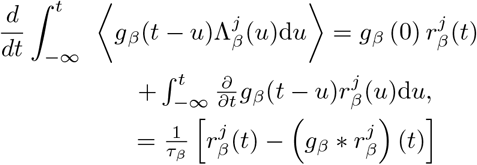

Starting from the last equality and writing 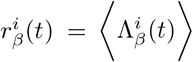, the Leibniz step with the normalized exponential kernel gives, term-by-term,

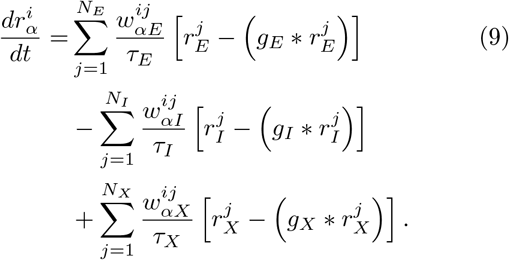

Averaging over population *α* yields:

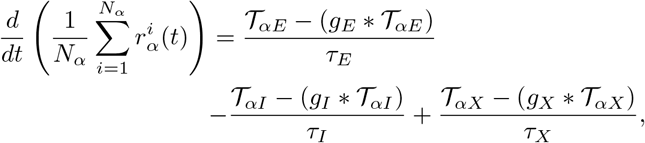

where the three sums have the form:

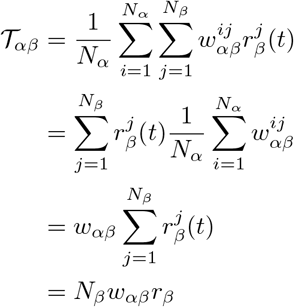

for *β ∈* {*E, I, X*}. Here we have assumed that the sum 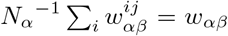 is independent of pre- and postsynaptic neurons, and we define the population-averaged firing rate 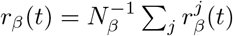.

Inserting the above into Eq. (9) we have:

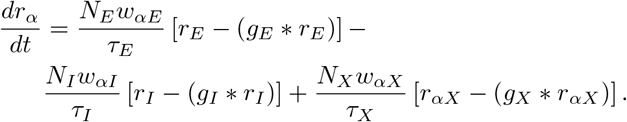

For the full *E* and *I* network, the evolution in vector form is:

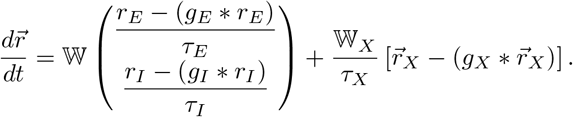

Here we have defined the matrices:

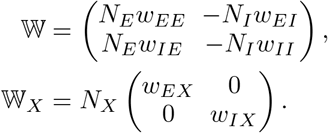

We can solve for 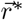 from the evolution equation above while retaining its initial constraint. For the normalized exponential kernels, we can recall the result of the Leibniz rule would still apply for the mean-field filtered activity

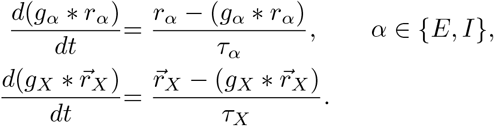

With the synaptic weights fixed, substituting these identities into the evolution equation gives:

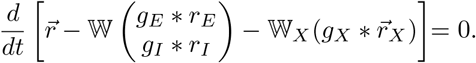

Integrating from 0 to *t* yields:

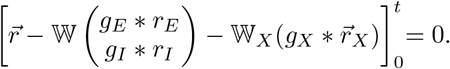

The initial rates and convolution histories satisfy the original mean-field integral equation, so the bracket at *t* = 0 is zero and remains zero for all time. For constant external input, a stationary solution satisfies 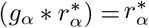 and 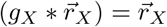, giving:

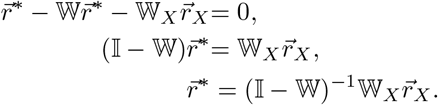

The final expression requires I − W to be invertible.

### B. Mean-field theory for synaptic dynamics

The STDP rule in Eq. (1) has a mean-field equivalent of the form:

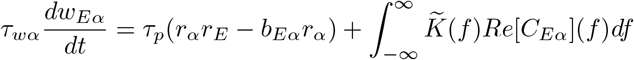

We define *C*_*Eα*_(*f*) = *S*_*α*_, *S*_*E*_ (*f*). For real stationary spike trains, *C*_*αE*_(*f*) = *C*_*Eα*_(−*f*) = *C*_*Eα*_(*f*)^*\**^, where * denotes complex coniugation. Further, take note that 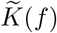 is the Fourier transform of the STDP kernel, which in this case is 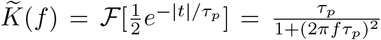. This kernel is integrated against the spike train cross spectrum ⟨*S*_*α*_, *S*_*E*_⟩ (*f*). For a network in the asynchronous state we have that ⟨*S*_*α*_, *S*_*E*_⟩ (*f*) ~ *O*(1*/N*). See [79] for the derivation.

### C. *E* − I network dynamics

#### 1. Stability of the Firing Rates

The dynamics of the fast *E* − *I* network are given by:

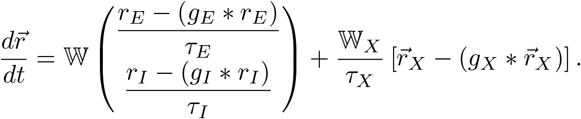

Holding the synaptic weights and external input fixed, the perturbations obey the following linearized equation, where *J* is a linear operator:

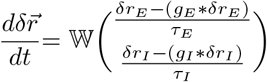

The mean-field integral equation of our system is:

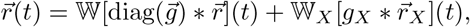

By inspection, we can quickly see this can be manipulated via differentiation to get:

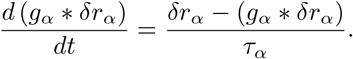

We see this gives us a relationship between the evolution of the perturbation, and the convolved perturbation:

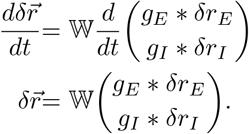

we can put the above into the linearized ODE

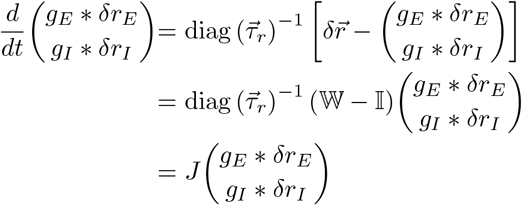

where,

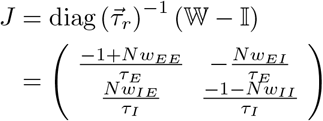

We can then obtain a quadratic characteristic equation whose roots are *λ*_*±*_.

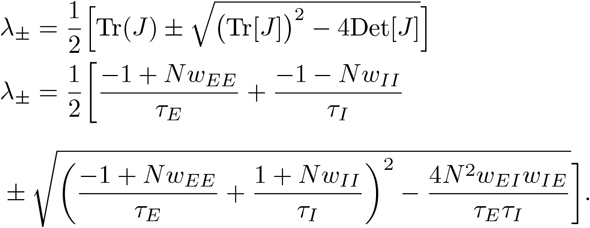

#### Case 1: Complex Case

The eigenvalues are complex when we have

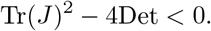

In this case, for the real component of the eigenvalues to be negative (to yield a stable solution), we then require

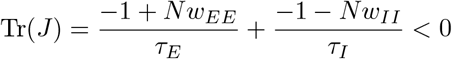

Rearranging gives the following condition:

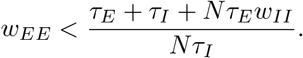

When the cross population weights are strong, then this is effectively an “oscillation” dominated regime in which the cross talk between populations is strong relative to the recurrent talk within populations. This produces an upper bound on the recurrent excitatory weights relative to the recurrent inhibitory weights. That is, when the cross talk is large, the recurrent inhibitory weights also need to be large in order for the firing rates to remain stable.

#### Case 2: Real Case

Recurrent weights large relative to cross weights all−→ real. In other words, this is the case in which the recurrent weights are strong relative to the cross-weights between the populations. The eigenvalues are real when

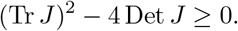

For both eigenvalues to be negative, their sum must be negative and their product positive: Tr *J <* 0, Det *J >* 0. In this case, the radical term is positive and the eigenvalues are purely real.

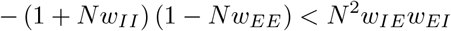

This condition creates an upper bound on the recurrent weights together.

#### Boundary Between Cases

It is clear that given these cases, there is also a boundary in phase space in which the imaginary (oscillatory) regime transitions to the real (non-oscillatory) regime with strong recurrent weights. This will occur when the radical term is zero in the solution for the eigenvalues above. If the term in the radical is negative, then we are in the imaginary oscillatory (strong cross *EI* weights) regime

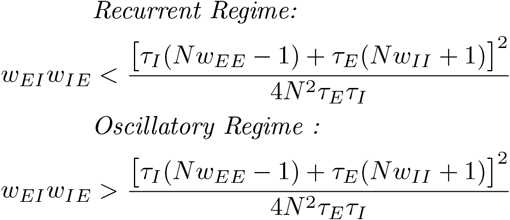

The boundary of the oscillatory regime defines a parabola in the *w*_*EI*_ vs *w*_*EE*_ plane, above which (large *w*_*EI*_ or *w*_*IE*_) the model is in the oscillatory regime and below which it is in the recurrent regime (small *w*_*EI*_ or *w*_*IE*_).

It is also worth calculating the boundaries in which the rates become negative. If 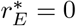, we see a boundary 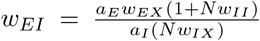 arises and if 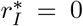, then the boundary is 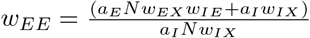.

#### 2. Fixed points of the Slow System: The Weights

We conduct a similar analysis as before, but now on the slow system in which we find the fixed points of the weights. In the asynchronous mean-field limit the correlation integral is *O*(1*/N*) so we neglect it, such that 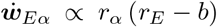. To study the steady state of the slow system, we similarly set 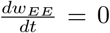 and 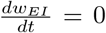, which occurs if *r*_*E*_ = *b* and *r*_*E*_ *>* 0. We assume that by the time the weights reach a steady state the rates are at their steady states such that 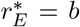. This is known as a fast-slow decomposition. This will produce the line attractor

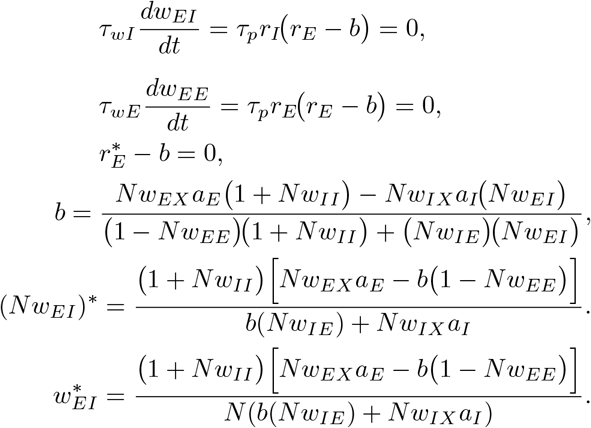

Recall we did all of this analysis because we assume a fast-slow decomposition in which the fast system reaches a steady state extremely rapidly relative to the slow system:

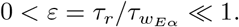

We can define a critical manifold *C*, which is the set of all points where the fast subsystem is already in equilibrium for the current weight values. This manifold exists in the absence of the weights changing.

We can consider the case of the weights evolving; as long as *ε* = *τ*_*r*_*/τ*_*w*_ ≪ 1, Fenichel’s theorem ensures the persistence of a locally invariant perturbed manifold *C*_*ε*_ that lies an *O*(*ε*) distance from the critical manifold *C*. The dynamics approach the line attractor in the (*w*_*EE*_, *w*_*EI*_) plane (the projection of *C*_*ε*_ onto weight space). Fenichel’s theorem relies on normal hyperbolicity of the fast subsystem equilibria (fast-direction eigenvalues have non-zero real parts) [123, 124].

In the model, 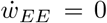 forces *r*_*E*_ = *b*. The dynamics approach the line attractor in the (*w*_*EE*_, *w*_*EI*_) plane, which can be considered a subset of *C*.

### D. Stability of Fixed Points for the Line Attractor

Our aim is to investigate stability in a recurrent system with excitatory and inhibitory plasticity. While this nonlinear *EI* plasticity rule has been shown to stabilize runaway excitatory weight dynamics, this has only been examined in feedforward models [37]. In order to study the role of *EE* and *EI* plasticity on network stability in the recurrent network, we use timescale separation between the rate and weight dynamics. We assume the firing rate dynamics occur more rapidly than the weight dynamics (*τ*_*wE*_, *τ*_*wI*_ ≫*τ*_*E*_, *τ*_*I*_), allowing us to analyze firing rate stability and weight stability separately. This reduces the four-dimensional problem to a two-dimensional problem. We will thus assume that the firing rates reach their fixed points before the plasticity dynamics become relevant. Here *h* = (*τ*_*p*_*/τ*_*wE*_)*r*_*E*_(*r*_*E*_− *b*) and *z* = (*τ*_*p*_*/τ*_*wI*_)*r*_*I*_(*r*_*E*_− *b*) are the mean synaptic weight velocities.

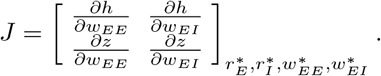

Using the fixed-point relation 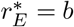, we get

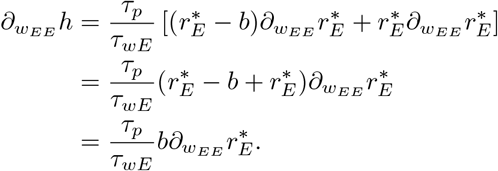

Likewise, the other terms of the Jacobian reduce to

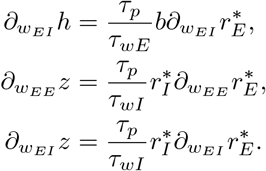

Thus,

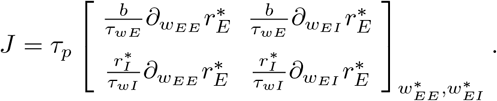

To test the stability of the line attractor, we leave *w*_EE_ as a free parameter and substitute the corresponding value of 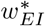, since the line attractor is a function of *w*_EE_. This gives

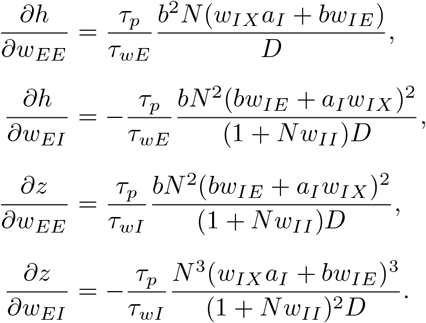

Here *D* = *w*_*IX*_*a*_*I*_(1 −*Nw*_*EE*_) + *Nw*_*EX*_*a*_*E*_*w*_*IE*_. To determine stability, we need to find the eigenvalues of the Jacobian. For a two-dimensional system,

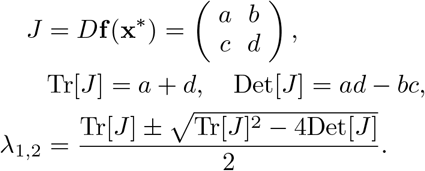

A direct inspection shows that the determinant of the Jacobian above is zero. Therefore, one eigenvalue is zero, corresponding to motion along the line attractor. Stability transverse to the line requires the nonzero eigenvalue to be negative. Since the nonzero eigenvalue is the trace, the stability condition is

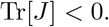

The trace is

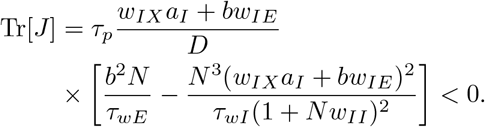

Since *w*_*IX*_*a*_*I*_ +*bw*_*IE*_ *>* 0, the sign of the trace is determined by D and the bracketed term. Ensuring the firing rates are stable and positive means D is positive. Therefore, we consider the bracketed term’s sign to determine the condition. Taking the bracketed term gives

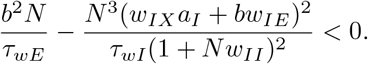

Multiplying through by the positive factors and simplifying gives

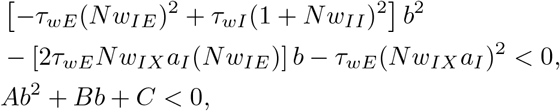

where

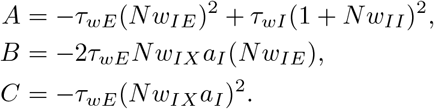

The discriminant is

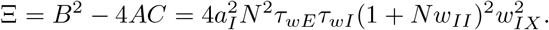

Because all parameters are positive, Ξ *>* 0. Thus, the quadratic always has two real roots. The remaining distinction is whether the parabola opens upward or downward, which is determined by the sign of *A*.

For *A >* 0,

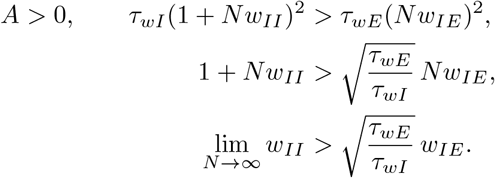

For *A <* 0,

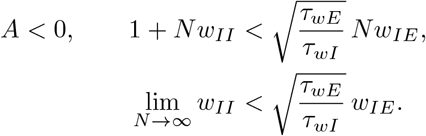

We now consider the two roots *b*_1_ and *b*_2_. The first root is

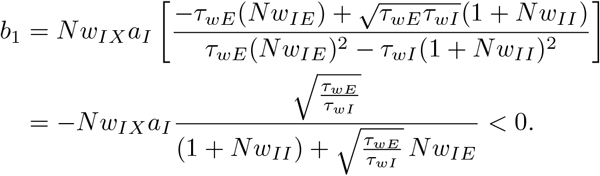

The second root is

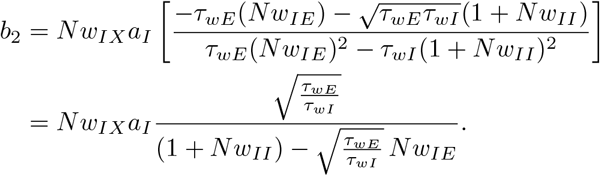

When *A >* 0, the denominator of *b*_2_ is positive and *b*_1_ is clearly negative. The part of the parabola that lies below the x axis thus falls between these two roots. Therefore, the viable b for the *A >* 0 case is,

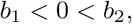

and the admissible interval is

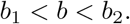

Thus, *b* is bounded above by *b*_2_.

When *A <* 0, the denominator of *b*_2_ is negative, so both roots are negative. In this case, the ordering is

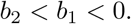

Likewise, the part of the parabola falling below the x axis will be to the left and right of these roots:

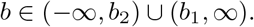

Since *b*_1_ *<* 0, any positive value of *b* lies in the interval (*b*_1_, ∞). Thus, in the *A <* 0 case, the quadratic condition does not impose an additional upper bound on positive *b*.

### E. Separatrix of the slow–weight dynamics

The slow weight dynamics are two-dimensional, so every trajectory lies in the (*w*_*EE*_, *w*_*EI*_) plane. In this plane, the line attractor is given by the set of weights satisfying 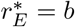. The region in which this attractor is rate-stable is bounded above by the 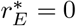 line and below by *w*_*EI*_ = 0 together with the rate-stability boundaries derived previously. Nevertheless, even for initial conditions within this nominally stable region, certain parameter choices yield trajectories that leave the physically relevant weight domain and do not converge to the line attractor. This motivates identifying additional invariant boundaries of the slow flow.

In particular, we seek a *separatrix* : an invariant curve that partitions weight space into regions with qualitatively different long-term outcomes (e.g., trajectories that approach the line attractor versus those that do not). Because the slow dynamics are deterministic and autonomous, trajectories cannot cross a separatrix; initial conditions on opposite sides evolve toward different asymptotic behaviors.

The separatrix of interest is the phase curve passing through a distinguished point in weight space where the steady-state rates satisfy 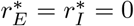. At this point, the closed-form fixed-point expressions *r*_*E*_ (*w*_*EE*_, *w*_*EI*_) and *r*_*I*_ (*w*_*EE*_, *w*_*EI*_) become singular (taking an indeterminate 0*/*0 form), so the reduced fast-slow description must be interpreted in a limiting sense. This point is therefore nonhyperbolic in the reduced slow flow and serves as the natural reference point for selecting the separatrix.

#### 1. Setup (fast–slow reduction and slow flow in weight space)

With rate dynamics fast relative to synaptic weights, the steady state rates are functions of the synaptic weights (*r*_*E*_(*w*_*EE*_, *w*_*EI*_), *r*_*I*_(*w*_*EE*_, *w*_*EI*_)):

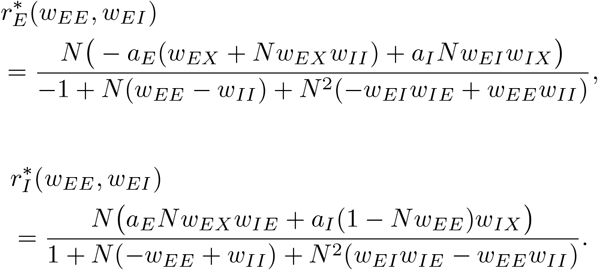

#### 2. Eliminate time and separate variables

We can obtain the trajectories by dividing out our differential equations. We will luckily be able to integrate these out to get actual trajectory curves in the (*w*_*EE*_, *w*_*EI*_)-plane. Dividing the slow equations (which removes the common factor 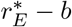) gives

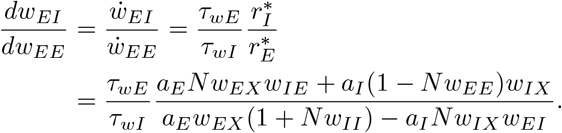

Separate variables and integrate:

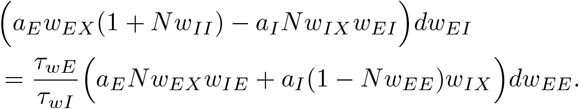

The integral that results is very simple:

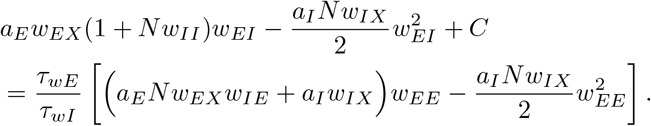

#### 3. How will this characterize the separatrices?

A separatrix is an invariant curve that forms the boundary between different trajectory behaviors. Thus the separatrix is the trajectory that passes through the reference (nonhyperbolic) fixed point. Thus, upon integrating, we will get the specific separatrices by using the coordinates of the nonhyperbolic fixed point to get the constant of integration. This point corresponds to 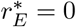 and 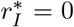, which in weight space has the following coordinates:

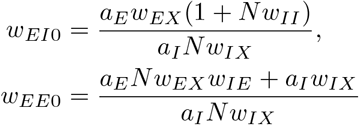

Evaluating the output of the integral at (*w*_*EE*0_, *w*_*EI*0_) gives the constant that selects the separatrix:

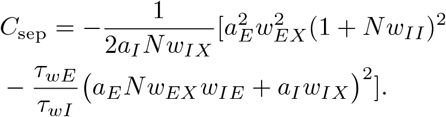

#### 4. Explicit separatrix w_EI_ (w_EE_) (two branches)

Substitute *C*_sep_ back in and solve the resulting quadratic in *w*_*EI*_. The separatrix (the phase curve through the fixed point) satisfies

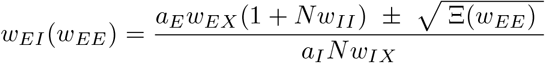

where the (nonnegative) discriminant simplifies to:

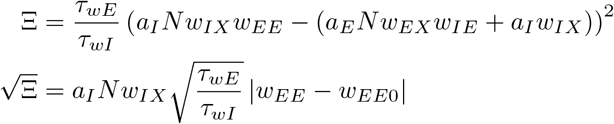

Plugging in for Ξ and then simplifying common terms further, this becomes very clean:

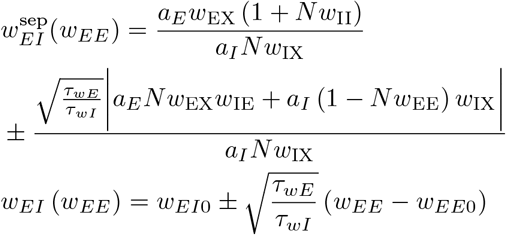

### F. Mean-field for the Triplet STDP Rule

The *E* →*E* plasticity rule considered in the main text (Eq. 1) is linear in the postsynaptic firing activity. If we consider the case in which the rule is quadratic in the postsynaptic neuron activity, which is the mean-field analog of the triplet STDP rule [83, 84], we see the same constraint on the plasticity threshold arises.

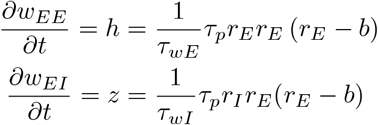

Keep in mind here, we use the fixed points of the firing rates from the fast system. The line attractor here 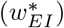 also remains unchanged due to the homeostatic rule. This will produce the same fixed point condition 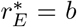, so we will see the same stability condition as in Appendix D:

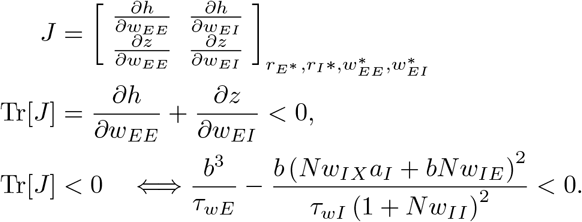

We see this is exactly the same result as what we have in the linear case. Thus the conditions for the linear and nonlinear plasticity rules are the same.

### G. Perturbations on the Line Attractor (Analytics)

This section helps build the mathematical relationship between the propagator Δ(*f*) and various response and stability properties of the network. As discussed, the propagator provides information about how the network responds to perturbations. Beginning with the integral equation of our mean-field model:

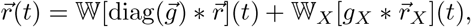

Observe that this is equivalent to the mean-field rate Eq.(2) if 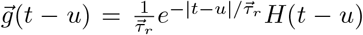. To perturb the system, we inject spikes directly into the *E* population.

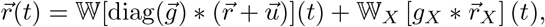

this drives fluctuations around the fixed point,

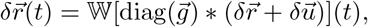

take into Fourier space

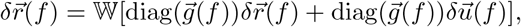

solve for 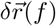,

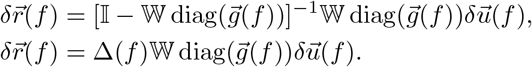

This analysis will allow us to relate the response to perturbations to the response via the propagator.

#### 1. Stability Boundaries from Propagator

Clearly the propagator plays a key role in determining how a response perturbation evolves due to a network perturbation. In Fourier space, this takes the form:

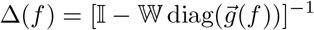

Given its relationship to the perturbation response, we can consider that the propagator can tell us where the boundaries of stability lie. These boundaries are determined when the denominator becomes singular. In particular, if det[I − W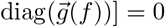] = 0, then the propagator has a pole (is undefined), meaning there exists a mode whose gain diverges. Equivalently, this happens when one eigenvalue of W diag 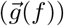 equals 1. Consider the denominator term:

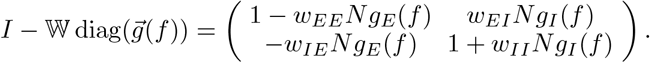

Here *g*_I_(*f*) and *g*_E_(*f*) correspond to the kernels with time constants *τ*_I_ and *τ*_E_ respectively. The determinant gives:

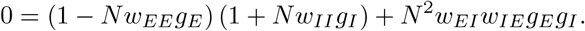

Substitute *g*_E_(*f*) and *g*_I_(*f*) :

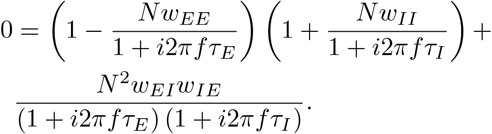

Multiply both sides by (1 + *i*2*πfτ*_E_) (1 + *i*2*πfτ*_I_) :

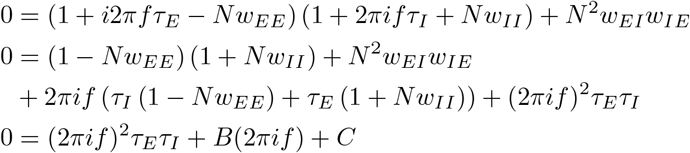

We can see this gives us a 2nd order polynomial in *z* = 2*πif*, with *A* = *τ*_E_*τ*_I_, *B* = *τ*_I_ (1− *Nw*_EE_)+*τ*_E_(1+*Nw*_II_), and *C* = (1− *Nw*_EE_)(1 + *Nw*_II_) + *N* ^2^*w*_EI_*w*_IE_. This allows us to invoke the Routh–Hurwitz criterion for stable systems. In particular, if we have our propagator which uses the first order kernel (e.g. *f*(*z*) = 1*/*(1 + *τz*)), then we can obtain the above quadratic characteristic polynomial

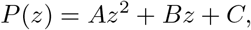

whose roots lie in the left half-plane if the system is stable. For a second-order system, stability is equivalent to

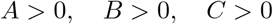

Considering *C >* 0 and *B >* 0 will correspond to conditions of stability for the non-oscillatory and the oscillatory mode respectively:

#### Non-oscillatory regime

If f=0, then the real part must vanish on the boundary

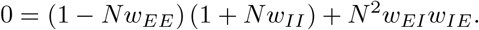

#### Oscillatory regime

We also consider when *f ≠* 0. In this case, both the real terms and the imaginary terms must vanish on the boundary.

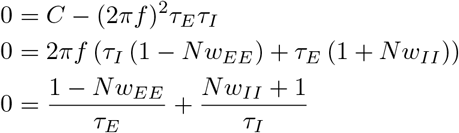

We also see B would need to be 0, but this would actually give the same equation!

Thus, this gives us equations for the boundaries! We can consider this will be stable for weak coupling *w* = 0 up to some strong coupling *w >>* 0, particularly for *w*_*EE*_, in which we can see that the equation in the DC case and the oscillatory case will be greater than 0. This can help us pinpoint an inequality in which we obtain stable dynamics, which will be 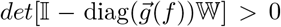. So we can simply say the two equalities are both positive inequalities, such that:

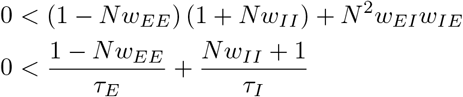

These are the same conditions of stability we would find were we to use the eigenvalues of the Jacobian.

#### 2. Dynamical Response to Perturbations

Let us begin with our expression of the perturbation in terms of the propagator. Write the (time-domain) evolution equation for 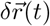:

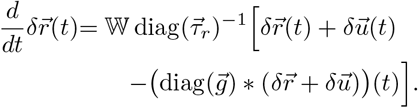

Here 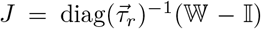 is the Jacobian of the filtered perturbations, rather than of 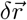 itself. Now consider rewriting this in terms of a function Φ(*t*), which is in terms of the propagator. It will give us the full evolution of the perturbation

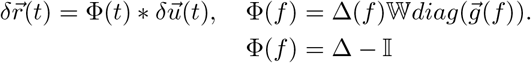

Using *g*_*α*_(*f*) = (1 + *i*2*πfτ*_*α*_)^*−*1^, we can rearrange the frequency-domain expression as:

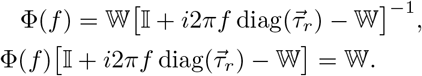

Expanding and multiplying from the right by 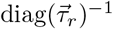 gives:

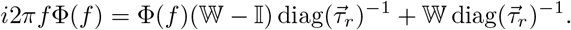

Taking the inverse Fourier transform and convolving with 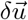 gives the following equations. Here 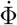 denotes the ordinary derivative for *t >* 0, extended causally, with the jump at *t* = 0 written separately:

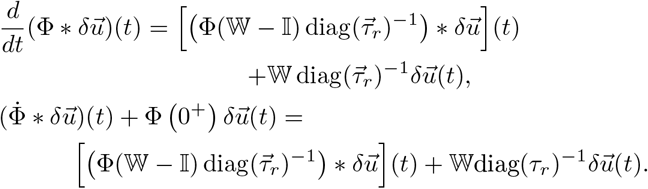

Rearranging gives:

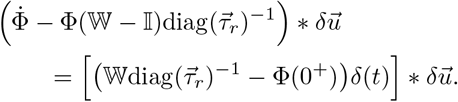

For this to hold for arbitrary 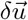, we must have: Φ (0^+^) = W diag(*τ*_*r*_)^*−*1^. This gives us a simple ODE for *t >* 0, for which we know the solution:

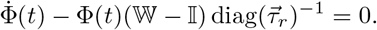

Using 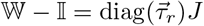, this is equivalently:

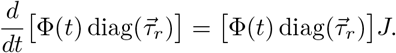

The solution to this equation can be written as the following Green’s function:

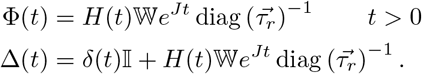

This will give us the evolution of the rates in response to a perturbation 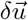:

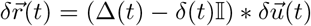

#### 3. Energy

We are interested in obtaining an expression of the energy in terms of the propagator (and network parameters). Consider an energy like term from the integrated perturbation, which can be written as follows: 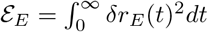.

We just need to solve for the perturbation, square it, and integrate it. The procedure is straightforward, beginning with our system of equations:

With synaptic weights held fixed and no perturbation of the external drive, the presynaptically filtered meanfield equations give

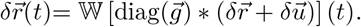

For invertible *W*, multiplying by *W*^*™*1^ and differentiating the exponential convolutions gives

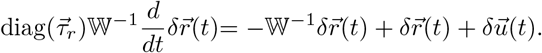

Using 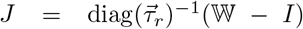, multiplying by *W* diag 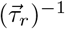 gives the following component equations for an E-only pulse:

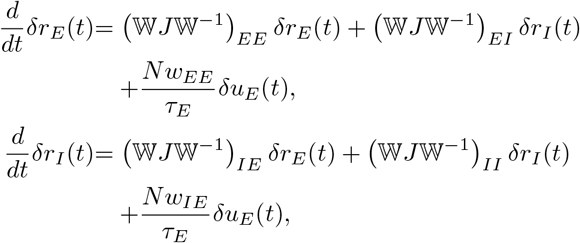

where the pulse injected only into E is, *δu*_*E*_(*t*) = *Aδ*(*t*), *δu*_*I*_(*t*) = 0. So for *t >* 0,

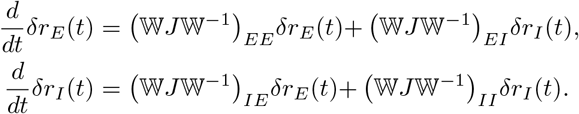

The invertibility assumption is used only for this intermediate calculation. The final expression for ℰ_*E*_ extends continuously to singular *W* whenever *J* remains stable.

We want to integrate these equations across time. Consider the initial conditions such that the left limits are 0 and the right limits are the jump due to the pulse: *δr*_*E*_(0^*™*^) = *δr*_*I*_(0^*™*^) = 0, and the delta-pulse gives the jump

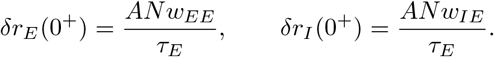

Because we know *δr*_*E*_(*t*) will be coupled with *δr*_*I*_(*t*), to get an equation for ℰ_*E*_, we will need the evolution equations of the squared perturbations of E, I and the mixed perturbations:

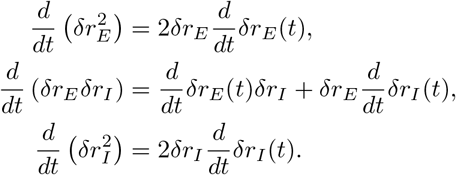

Plugging in for 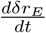 and 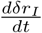 we can get:

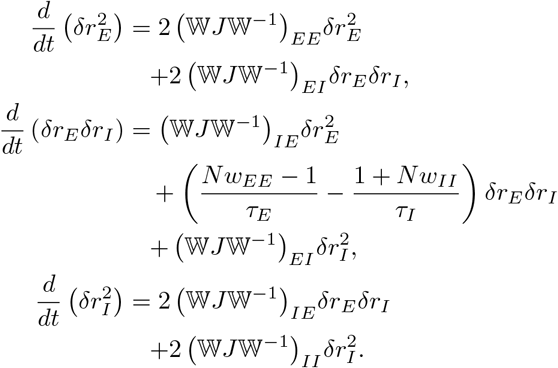

The coefficient of *δr*_*E*_*δr*_*I*_ in the mixed-product equation is unchanged because tr(*W J W*^*™*1^) = tr *J*.

Now integrate each from 0^+^ to ∞. Since the fixed point is stable, both perturbations decay to zero as *t* → ∞. That gives

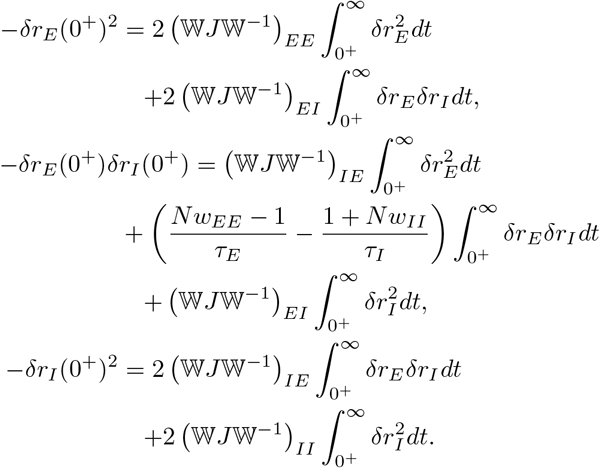

This is a linear system for the three unknown integrals

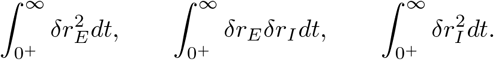

Solving that system for the first one gives:

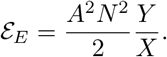

where

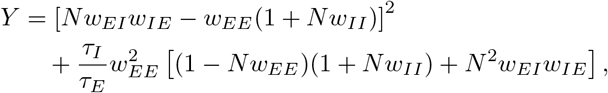

and

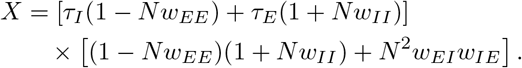

Getting this in terms of the propagator, we can consider:

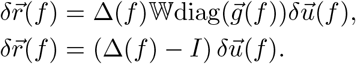

Given that

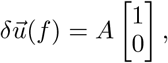

we see that *δr*_*E*_(*f*) = *A* (Δ_*EE*_(*f*) ™ 1).

Parseval’s Theorem therefore gives us the energy in terms of the propagator in the frequency domain.

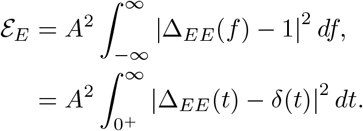

Some tedious algebra can likewise show that this can be further written in terms of the propagator:

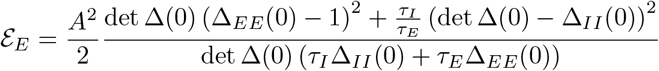

#### 4. Relating Propagator to Jacobian

Likewise, if we want an expression of the propagator in terms of the more familiar Jacobian, we can manipulate the expression a little more:

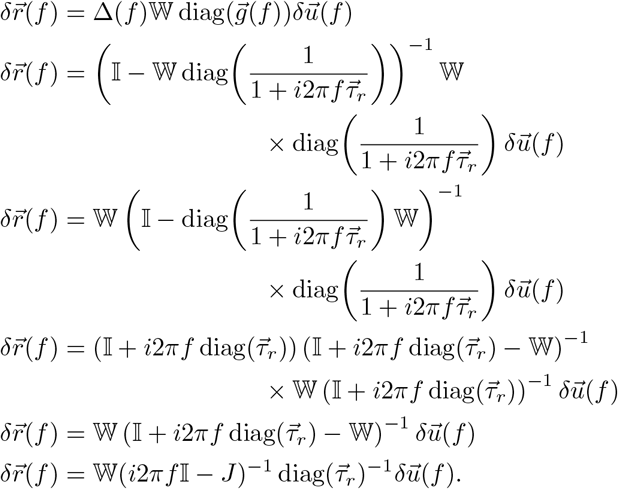

This allows us to quickly see that:

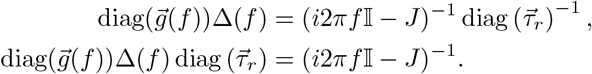

#### 5. Relaxation Time from the Propagator

This section is meant to show how to relate the eigenvalues of the propagator to the relaxation time of the system in response to a perturbation. We will begin by considering that typically relaxation timescales of a dynamical system are obtained from the eigenvalues of the Jacobian. Specifically, in response to a perturbation, the evolution of such a perturbation is represented through the Jacobian :

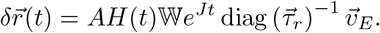

If we feed a rapid delta function perturbation, because *W* changes only on the plasticity time-scale, *J* is effectively frozen while the *δ*-step is active. If *J* is diagonalisable, *J* = *V* Λ*V* ^*™*1^ with Λ = diag (*λ*_+_, *λ*_*™*_), then

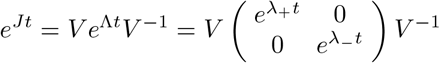

Here V is the matrix of column eigenvectors. Let *λ*_+_, *λ*_*™*_ be the eigenvalues of *J* (complex if oscillatory). The two eigenvalues *λ*_+_, *λ*_*™*_of *J* tell one how small perturbations decay (or grow) along two independent directions (the eigenvectors).

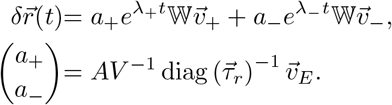

The slow time scale of the perturbation is thus proportional to 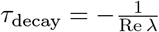. The eigenvalue whose real part is closest to zero will correspond to the slowest timescale: For a stable system, the eigenvalue whose real part is closest to zero determines the slow relaxation timescale:

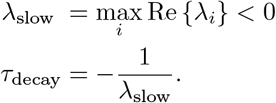

For our system, when the eigenvalues form a complexconjugate pair we merely need to find the real component of a given eigenvalue. This gives us the timescale from the Jacobian itself, but in the subsequent paragraphs, we show how to relate this timescale to the eigenvalues of the propagator Δ(*f*).

We can first consider the eigenvalue equation of the Jacobian and manipulate it until we have *J* written directly in terms of the zero-frequency propagator. This is because we can recall we have a relationship between the propagator and the Jacobian:

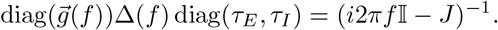

Therefore, if we evaluate this expression at zero frequency and invert it, we can recover the eigenvalues of the Jacobian from the propagator. Evaluating at *f* = 0 gives

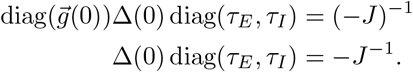

Now invert both sides:

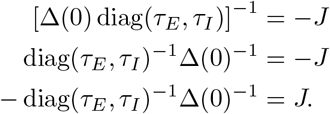

Now, we can consider an eigenpair of the Jacobian first:

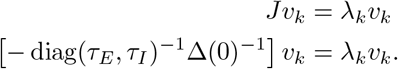

Thus, the Jacobian eigenvalues can be written in terms of the propagator as

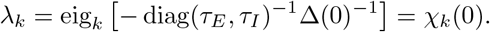

We can then manipulate these eigenvalues to be in terms of the decay time:

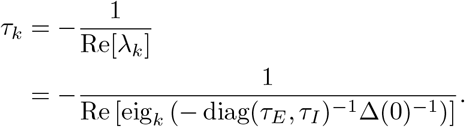

The slow relaxation time is set by the eigenvalue whose real part is closest to zero. Therefore,

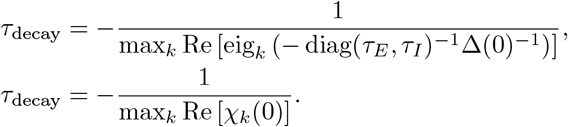

#### 6. Trial-to-trial variability

Now, to model trial-to-trial variability, we consider the mean-field intensity cross-spectrum:

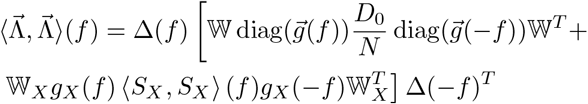

We evaluate the inverse Fourier transform of the cross spectrum at 0 lag, which is just the integrated cross spectrum. We plot the first entry of this 2 × 2 matrix along the line attractor in **Fig.5i**. We clearly see the propagator plays a defining role in shaping the cross spectrum, and thus the trial-to-trial variability. Appendix G 7 derives the full-dimensional expression; Appendix G 8 derives the mean-field expression.

#### 7. Covariance of the Hawkes process

One advantage of modeling neural networks with Hawkes processes is the fact that we have a closed form expression for the covariance [71, 72]. The full dimensional Hawkes system obeys:

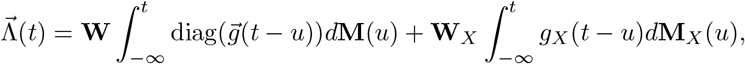

where we denote the spike train from neuron *j* of population *α* as 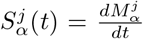. We can likewise incorporate this spiking notation directly into the convolution:

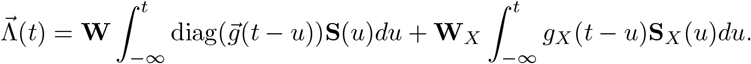

We relate the intensity of the Hawkes process to a realization of the spiking activity via:

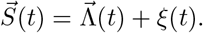

The statistics of *ξ*(*t*) carry the shot noise information of the Hawkes process, and satisfy ⟨*ξ*(*t*)⟩ = 0 and 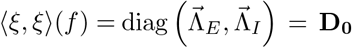. The second moment comes from the fact that Hawkes processes are conditionally independent Poisson processes.

The Fourier transform of 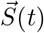 is simply:

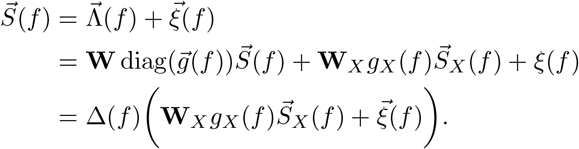

The last line comes from isolating for 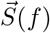 and writing 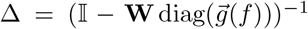. This yields the following expression for the cross-spectrum of 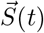

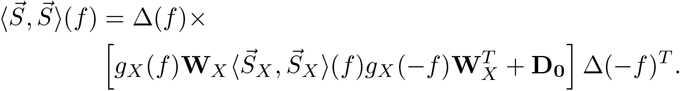

Likewise, using the definition of the intensity function in Fourier space, we see that:

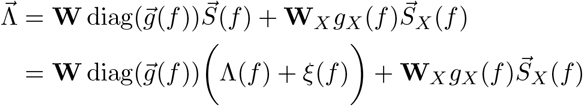

we can write the intensity function as:

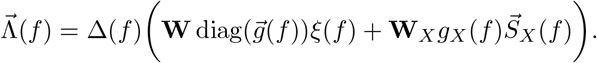

This gives the cross-spectrum of 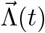 as:

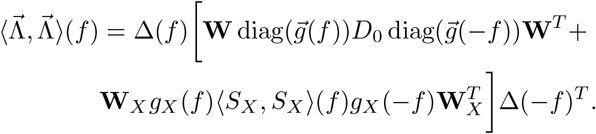

This subtle difference will allow us to consider how the spike train correlations impact the evolution of the plasticity rule (⟨*S, S*⟩) and the trial-to-trial variability of the mean firing rate (⟨Λ, Λ⟩). Note, this section is the only place where the propagator Δ is defined in the full dimensional form 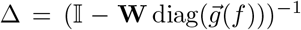. In subsequent sections, Δ will refer to the 2 dimensional object in the mean-field picture, namely, 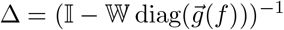.

#### 8. Mean-field Covariance

We will derive the mean-field spike train cross spectrum of the recurrent Hawkes network here. This will be useful for the mean-field plasticity rule. We will first take the summed activities of all of the neurons in each population. This will effectively reduce the system to a 2D system. The sum of the Hawkes processes is still a Hawkes process, so we can use the general form of the Hawkes covariance on the 2-dimensional system to obtain the cross spectrum on the summed activities. Then, to obtain the average covariance, we divide by *N* ^2^ to get the average covariance between neurons.

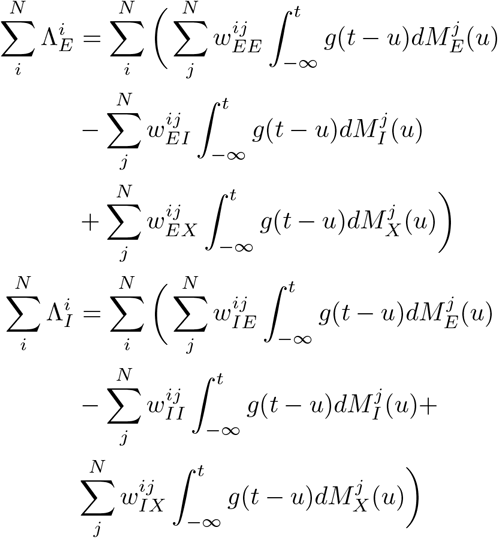

We treat each neuron as effectively identical (certainly statistically) and use 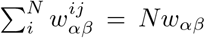, where *w*_*αβ*_ is the mean connection from population *β* to population *α*. We also assume the synaptic filtering kernels *g*(*t* ™*u*) are identical within a population.

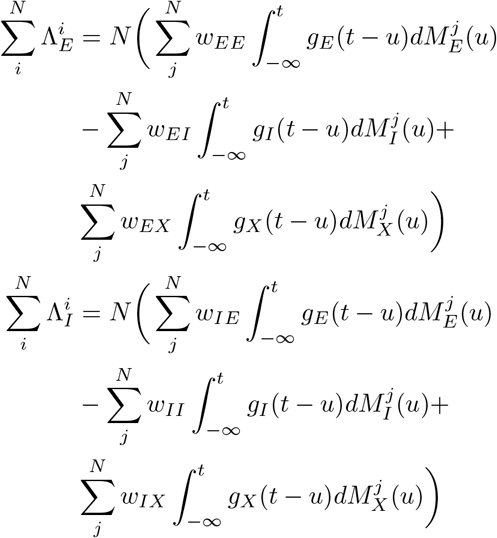

We can move the sums over j into the integrals

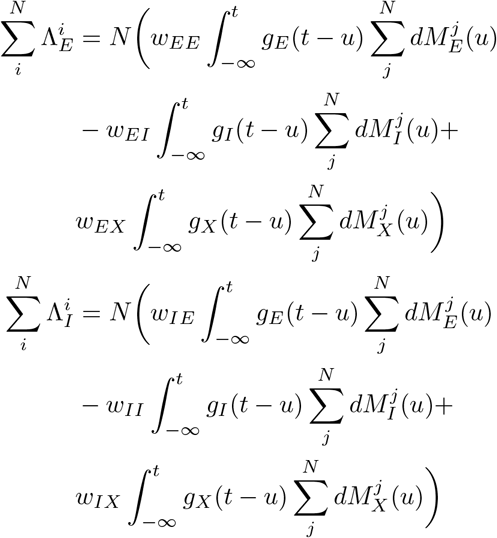

The above is now effectively a 2 dimensional system of the summed activity.

The goal is to compute the spike train cross spectrum of the 2-dimensional system 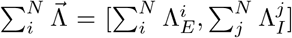, which consists of the total activities of the two populations. We can introduce the notation here that 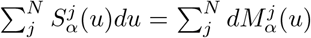. The cross spectrum of the summed process is similar to that seen in the previous section:

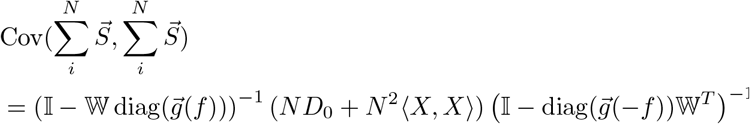

Here, 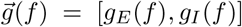. 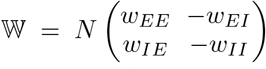. ⟨*X, X*⟩ is the mean-field cross spectrum of the external population, which is the 2 × 2 matrix. We will calculate this further below. Note though, these terms will correspond to the off-diagonals for the full *N* × *N* matrix, so we would expect to have *N*(*N* ™ 1) terms, which we will take to be roughly *N* ^2^ for large enough N. *D*_0_ = *diag*[Λ_*E*_, Λ_*I*_] is the diagonal matrix of the mean activity of a neuron, of which we have N diagonal entries. This is the spike train cross spectrum of the summed Hawkes processes, which only relies on the linearity of the Hawkes system. Since we are interested in the average covariance per neuron, we divide by *N* ^2^ to obtain:

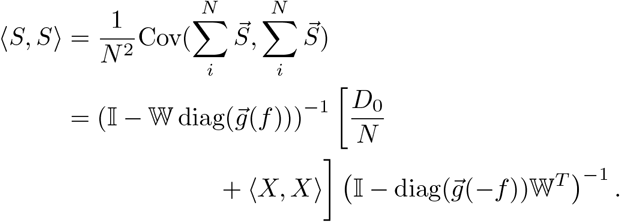

Here 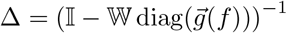, which is known as the propagator. 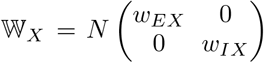. 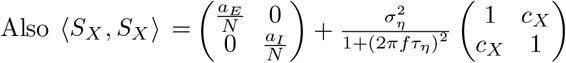.

Now we are almost there. We need to further determine what ⟨*X, X*⟩ looks like in the mean-field picture. Consider first that the total filtered activity from the connecting external neurons for a given recurrent neuron i in population *α* can be written as: 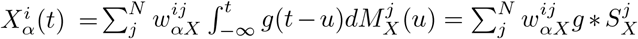, but we are going to derive the cross spectrum, which will entail writing 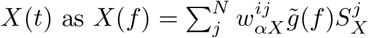

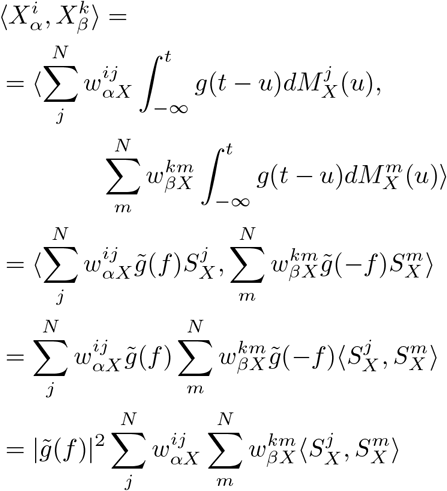

Here 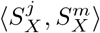 is the mean-field spike train cross spectrum of the external populations, which in this case are just inhomogeneous Poisson processes. The intensity functions for the respective populations are 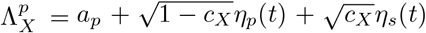, where *p* ∈ *E, I*. Here *η*_*s*_(*t*) and *η*_*p*_(*t*) are Ornstein Uhlenbeck noise processes.

We can split the final line into the auto-spectrum terms (of which we will only have a single sum representing the diagonal) and the cross spectrum terms (of which we will have N(N-1) terms).

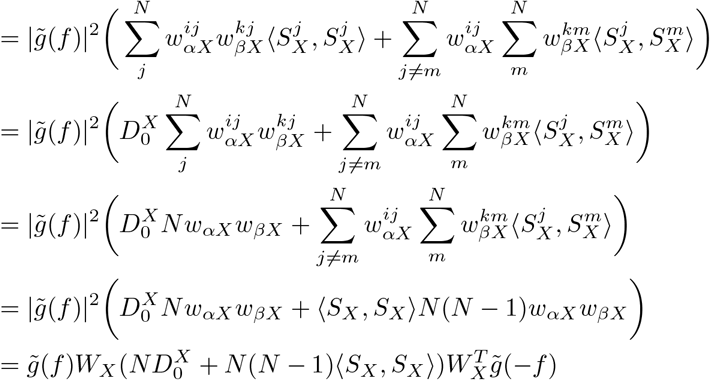

Here the auto-spectrum is the diagonal of the external neural activities - which for Poisson neurons is just the mean firing rate such that 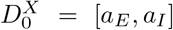. Further, 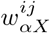 and 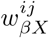 have been replaced with their mean values ⟨*w*^*ij*^⟩ = *w*_*αX*_ where *w*_*αX*_ is the mean connection from population *X* to *α*. Because we use weights selected from a Gaussian distribution, this would hold for a large network. Here ⟨*S*_*X*_, *S*_*X*_⟩ is the average covariance between external populations. Because we have two external populations, this is a 2 × 2 matrix such that 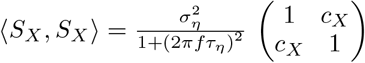.

This is if *σ*_*η*_ = *σ*_*s*_ for the OU processes. If they are different then 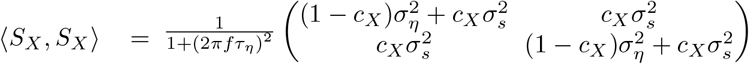.

## Notes

### Competing Interest Statement

The authors have declared no competing interest.

### Summary of Updates

We have corrected small mistakes including some notational corrections in the math in the main text and in the appendix including figure 2, figure 5.

